# Cortical myelination networks reflect neuronal gene expression and track adolescent age in marmosets

**DOI:** 10.1101/2025.09.26.678906

**Authors:** E D Hutchings, S J Sawiak, R A I Bethlehem, A C Roberts, E T Bullmore

**Author notes:** Correspondence to E D Hutchings.

## Abstract

Structural similarity provides a powerful framework for measuring coordinated macro- and microstructural variation across the cortex of a single brain. Similarity networks derived from myelin-sensitive MRI sequences undergo marked reorganisation during adolescence, linked to psychosocial outcomes in humans and rodents. However, the cellular mechanisms of myelination similarity and its development in non-human primate cortex remain unexplored. Here, we used myelin-sensitive T1w/T2w ratio images from a cross-sectional sample of 446 common marmosets (aged 0.62 to 12.75 years) to estimate cortical similarity networks in individual animals. Cortical areas with similar myeloarchitecture showed highly similar patterns of gene co-expression in glutamatergic neurons and PV+ and VIP+ interneurons, reflecting the activity dependence of myelination. A reliable age-based signal exists within network features, with coordinated developmental trajectories observed across the cortical hierarchy from primary to transmodal association cortices - a pattern that mirrors findings in human cortex. Taken together, marmosets demonstrate phylogenetically conserved patterns of myelination network development, potentially underpinned by key neuronal cell types that shape the functional specialisation of cortical areas.

## Introduction

Network science has significantly advanced our understanding of brain structure (Bullmore and Sporns 2009; Bassett and Sporns 2017). Magnetic resonance imaging (MRI) has been a key driver of this progress because it has enabled the collection of whole brain images at the millimetre scale in vivo (Bullmore and Sporns 2009). Analysis of MRI images allows quantification of brain structure both at the macro- scale, through metrics such as cortical thickness, surface area, and mean curvature; and at the micro-scale, below the minimal resolution of single voxels, using sequences such as diffusion imaging and magnetisation transfer (Lerch et al. 2017). Recent technical advances in structural similarity have permitted the integration of these MRI-derived structural features into a single network representation, with edges between nodes defined by the statistical relationship between the MRI features measured in pairs of regional nodes (J. Wang and He 2024; Sebenius, Dorfschmidt et al. 2025). Similarity networks therefore represent spatially patterned structural variation across the cortex (the architectome), rather than *directly* measuring connectivity (the connectome), as is the case in diffusion tensor imaging (DTI) tractography. However, areas that are structurally similar are more likely to be axonally connected due to the network formation principle of homophilic attachment, or “like attracts like” (Sebenius, Dorfschmidt et al. 2025). Accordingly, the architectome represented by MRI-derived structural similarity networks has been correlated empirically with the connectome represented by axonal tract tracing data in the macaque (Sebenius et al. 2023; Seidlitz et al. 2018) and rat (Smith et al. 2024) cortex.

A recently developed method for estimating structural similarity networks from MRI data is Morphometric Inverse Divergence (MIND), which uses Kullback-Leibler divergence to quantify how similar a pair of brain regions is in terms of their voxel- or vertex-wise distributions of one or more structural features (Sebenius et al. 2023). MIND and adjacent methods are technically robust and reliable (H. Wang et al. 2016; Sebenius et al. 2023), heritable (Sebenius et al. 2023), associated with similarity in gene expression (Sebenius et al. 2023; Seidlitz et al. 2018), developmentally sensitive (Sebenius et al. 2023; J. Wang and He 2024; Smith et al. 2024), and altered in psychiatric disorders (J. Wang and He 2024; García-San-Martín et al. 2025).

Animal models enhance our understanding of the biological underpinnings of structural similarity networks and how they are reorganised during development and disease. The common marmoset (*Callithrix jacchus*), a New World primate, represents a key rung on the translational ladder between rodent and human neuroscience, combining the tractability of rodent research with the translational relevance of primate research (Okano et al. 2012). Like rodents, marmosets have an accelerated lifespan compared to humans, reaching adulthood at 18-20 months (Schultz-Darken, Braun, and Emborg 2016; Sawiak et al. 2018), which facilitates longitudinal MRI studies that collect repeat measures over the lifespan. Like humans, marmosets show prolonged postnatal development in the context of familial nurturing (Burkart and Finkenwirth 2015; Miller et al. 2016), and protracted development of association cortex (Sawiak et al. 2018), particularly those areas involved in social cognition (Cerrito et al. 2024).

Representations of the marmoset structural connectome generated from “gold standard” tract tracing data (Majka et al. 2020; Watakabe et al. 2023) have revealed topological similarities to the human brain, with a rich club of highly interconnected hubs and exponentially distributed connection distances or wiring costs (Liu, Zheng, and Misic 2020; Theodoni et al. 2021). These topological parallels between human and marmoset brain networks, combined with the suitability of marmosets for MRI (Schaeffer et al. 2020) and the drive towards increasing non-invasive imaging of primates as part of the 3Rs in neuroscience (Prescott and Poirier 2021), have motivated efforts to generate MRI structural networks in the common marmoset, using DTI tractography (Hata et al. 2023) and structural covariance (Quah et al. 2022). However, there are problems with both of these approaches. DTI tractography suffers from a high false positive rate, i.e., identifying tracts between cortical areas that are not validated by comparable tract tracing data (Maier-Hein et al. 2017); and a bias against long-range connections, i.e., failing to identify tracts between physically separated cortical areas that are demonstrated by tract tracing data (Donahue et al. 2016). Whereas, structural covariance network analysis is constrained to estimate a network representing between-subject covariance, typically from a dataset of N∼100 single feature MRI scans, and therefore cannot provide the single-brain network analysis that is required for MRI studies of brain network development. We anticipated that similarity network analysis of MRI data in the common marmoset could circumvent these issues and provide a robust methodological platform for measuring the neurodevelopmental trajectories of primate brain network maturation.

To investigate the biological significance of structural similarity network analysis in the common marmoset, we focused on a myelin-sensitive MRI parameter, the T1w/T2w ratio (Glasser and Van Essen 2011). Myelin is important to normative brain function (Nave and Werner 2014), shows characteristic patterns of variation across the cortex (Burt et al. 2018; García-Cabezas, Hacker, and Zikopoulos 2020), and a clear programme of developmental changes in human (Flechsig 1901, Yakovlev and Lecours 1967). However, developmental changes in myeloarchitecture of the marmoset cortex have not previously been investigated. To do this, we estimated MIND networks from an open cross-sectional dataset containing T1-weighted (T1w) and T2-weighted (T2w) MRI images from 446 animals spanning pre-puberty to senescence (0.62 to 12.75 years; Hata et al. 2023). Specifically, we estimated the ratio between T1w and T2w images to yield parameter maps with enhanced myelin-sensitive contrast (Glasser and Van Essen 2011) and estimated similarity networks from parcellated voxel-wise T1w/T2w ratio images using MIND.

We used these MRI-based network estimates to test two key hypotheses grounded in gene expression and development of the marmoset brain. First, assuming that T1w/T2w MIND network metrics are representative at macro scale of cortical (co-)variation in microscopic myeloarchitectonics, they should be significantly co-located with molecular and cellular markers of myeloarchitecture. To test this, we benchmarked T1w/T2w MIND networks against two transcriptomic datasets: (i) spatially-resolved in-situ hybridisation images of myelin basic protein (*MBP*) messenger RNA (mRNA) in an infant and adult marmoset (Shimogori et al. 2018; Kita et al. 2021); and (ii) whole genome single nucleus RNA sequencing data from 12 cortical regions in 6 adult marmosets (Krienen et al. 2023). Second, assuming that T1w/T2w MIND networks can be validated as cortical myelination networks, we predicted: (i) that T1w/T2w MIND networks would show age-related changes during adolescence which are consistent across individuals and predictive of chronological age; and (ii) that the cortical patterning of network changes is conserved across species, such that cortical areas occupying a similar position on the hierarchy from low-level primary sensory areas to higher-order transmodal association areas (Mesulam 1998; Sydnor et al. 2021) would show similar trajectories of myelination network development, as has been observed in human (Paquola et al. 2019).

## Results

### Estimation and anatomical validation of T1w/T2w similarity networks

We estimated cortical T1w/T2w ratio (henceforth T1w/T2w) similarity networks in 446 individual marmosets aged between 0.62 and 12.75 years (Hata et al. 2023) using Morphometric Inverse Divergence (MIND; Sebenius et al. 2023; **Figure 1A**, **Supplementary Figure 1**). MIND networks use Kullback-Leibler (KL) divergence (Kullback and Leibler 1951) to quantify how similar or convergent every pair of cortical areas is in terms of their voxel-wise T1w/T2w distributions. Several methods have been used to estimate KL divergence in the context of MRI structural similarity networks (**Supplementary Methods**), though these methods have not been formally compared. We therefore compared two estimators KL divergence, based on either global or local comparisons of distribution density, and variable parameters of each of these estimators (**Supplementary Table 2, Supplementary Figure 2-3**), on a set of biologically informed benchmarks and an age prediction task. While both algorithms produced similar networks, the local estimator out-performed the global estimator across all benchmarks and age prediction (**Supplementary Figure 4**). Our downstream analyses therefore used the local KL estimator, as previously implemented in MIND (Sebenius et al. 2023), to construct similarity networks from voxel-level data.

**Figure 1:**
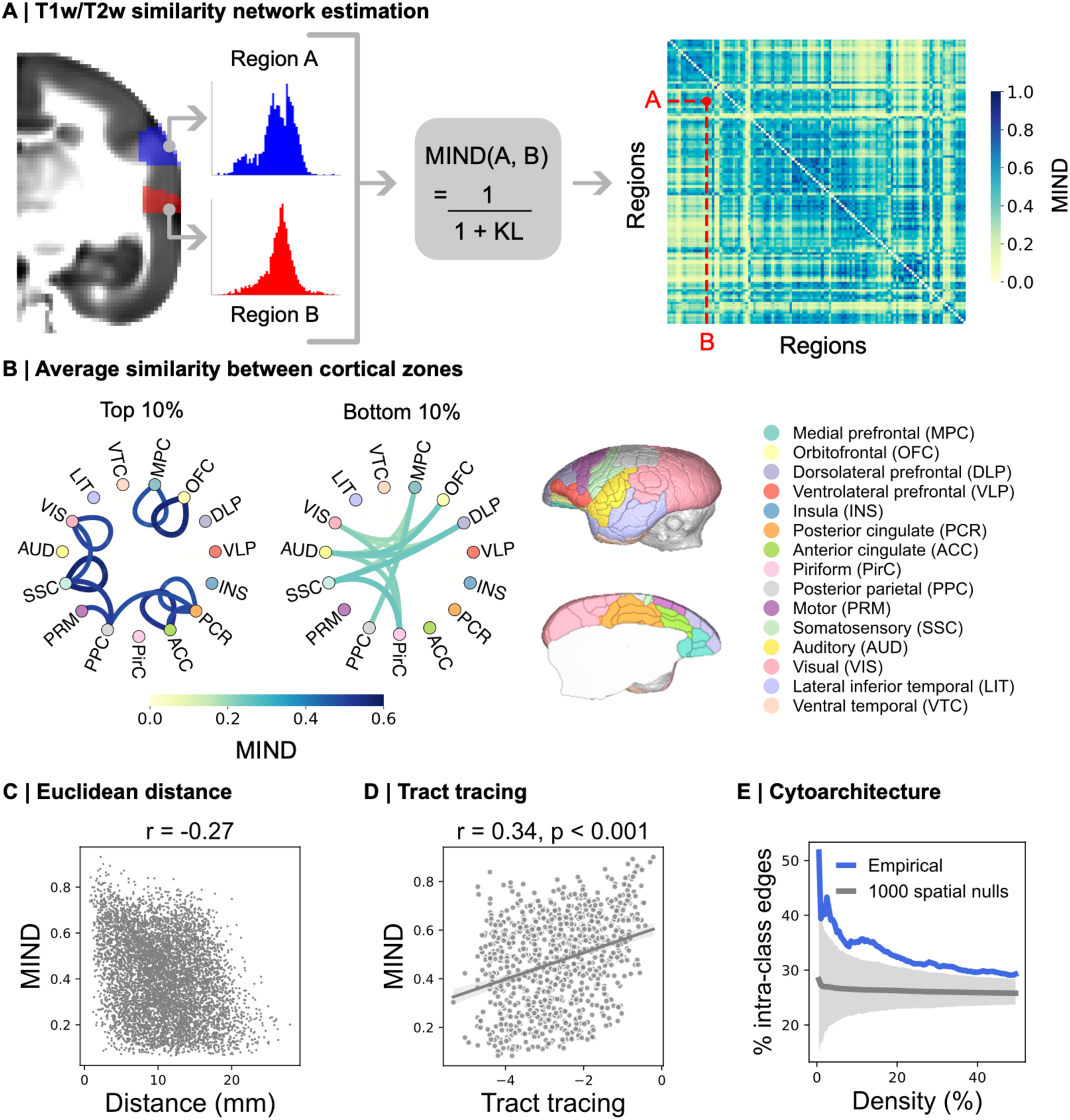
T1w/T2w similarity reflects axonal connectivity and cytoarchitectonic similarity. **A** Similarity between pairs of cortical areal T1w/T2w voxel distributions was estimated using Morphometric Inverse Divergence (MIND; Sebenius et al. 2023). The average adult T1w/T2w similarity network is shown on the right of the panel. **B** Mean similarity between cortical zones. Top and bottom 10% of mean zone-zone similarity is shown. **C** T1w/T2w similarity decays with distance. **D** T1w/T2w similarity is significantly correlated with axonal connectivity. **E** The proportion of edges between regions of the same cytoarchitectonic class increases as the network is thresholded to lower densities. The empirical network is shown in blue. The mean of 1000 spatial autocorrelation-preserving null networks is shown in grey, with shading indicating the 95% confidence interval.

The average adult (N=340, mean age = 4.3 years) T1w/T2w similarity network estimated using MIND displayed high average similarity within anatomically and functionally-defined cortical zones (Paxinos et al. 2012) and strong divergence between primary sensory and prefrontal association cortical zones (**Figure 1B**). Furthermore, the average adult network recapitulated biological relationships with Euclidean distance, axonal connectivity, and cytoarchitecture at the edge level shown to be present in structural similarity networks in other non-human species (Seidlitz et al. 2018; Sebenius et al. 2023; Smith et al. 2024). T1w/T2w similarity was anti-correlated with the Euclidean distance between regions (Spearman r = -0.27, **Figure 1C**). Using retrograde tract tracing data as a “gold standard” estimate of the average adult marmoset connectome (Majka et al. 2020), we observed a significant positive correlation between T1w/T2w similarity and axonal connectivity (Spearman r = 0.34, p_variogram_ < 0.001, **Figure 1D**), reflecting homophily in cortical connectivity (Sebenius, Dorfschmidt et al. 2025). Using a cytoarchitectonic class assignment map from (Atapour et al. 2024) (**Supplementary Figure 5B**), the proportion of edges between regions of the same cortical type increased as the network was thresholded to progressively lower densities, remaining above the average of 1000 spatially constrained null models (**Figure 1E**). This suggests that high values of T1w/T2w MIND represent the high cytoarchitectonic and myeloarchitectonic similarity between cortical areas of the same histologically defined class.

### T1w/T2w similarity networks align with molecular and cellular transcriptomic markers of myeloarchitecture

Owing to the strong correlation of T1w/T2w ratio MRI with myelin (Glasser and Van Essen 2011; Glasser et al. 2014), we hypothesised that T1w/T2w similarity would mirror similarity in a molecular transcriptomic marker of myeloarchitecture. To test this hypothesis, we downloaded spatially-resolved images of myelin basic protein (*MBP*) messenger RNA (mRNA) expression from the Marmoset Gene Atlas (Shimogori et al. 2018; Kita et al. 2021), measured using in-situ hybridisation, and warped these images to MRI space (**Supplementary Figure 6**). The resulting *MBP* mRNA volume was compared with an average adult T1w/T2w volume, formed by averaging N=340 spatially co-registered T1w/T2w scans with a mean age of 4.3 years. Both modalities show a drop-off in myelin-related signal in superficial layers of cortex, though this is more obvious in the *MBP* mRNA image (**Figure 2A**). Regionally parcellated cortical maps of mean *MBP* expression and mean T1w/T2w (**Figure 2B**) were positively correlated (Spearman r = 0.68, p_variogram_ < 0.001, **Figure 2C**). Standard deviation and skewness, quantifying the shape of each regional distribution, were less strongly, though still significantly, correlated between *MBP* mRNA and T1w/T2w MRI images (**Supplementary Figure 7**).

**Figure 2:**
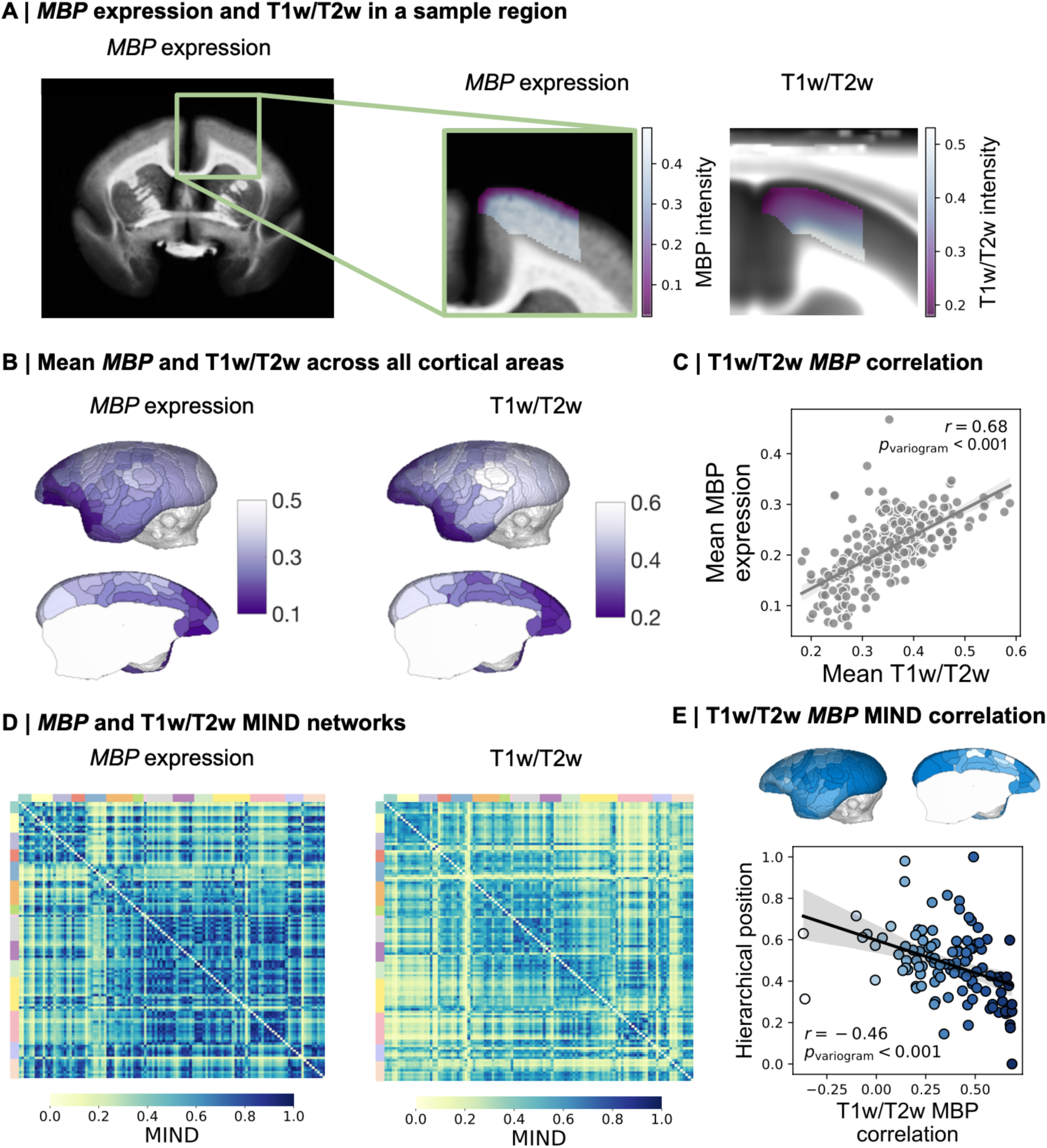
T1w/T2w similarity aligns with inter-areal similarity of myelin basic protein (MBP) gene expression. **A** Qualitative comparison of cortical myelination in MBP mRNA and T1w/T2w images in an example region, the left primary motor cortex. For T1w/T2w, the colour scale is capped at the 90th percentile of intracortical values to improve visualisation. **B** Mean MBP expression and T1w/T2w per cortical area. **C** Regional mean MBP expression and T1w/T2w were significantly correlated across all cortical areas. **D** Adult MIND networks of inter-areal similarity of MBP expression and T1w/T2w, grouped into cortical zones. **E** Regional correlations between the MIND network edge weights linking each cortical area to the rest of the brain in MBP and T1w/T2w MIND networks. Spearman’s rho for the correlation between MBP MIND and T1w/T2w MIND edge weights is shown for each area on a cortical map (top panel). The strength of positive correlation was greater for specialised sensory and motor areas positioned lower in the cortical hierarchy (bottom panel).

Given the high correspondence between T1w/T2w and *MBP* expression, we further hypothesised that *MBP* MIND networks would show a highly similar organisation to T1w/T2w MIND networks. Since each cortical area in the *MBP* mRNA image contains a distribution of voxel values of *MBP* expression, we could estimate the *MBP* MIND network using identical methods and a commensurate cortical parcellation as was used to estimate T1w/T2w MIND networks (**Figure 2D**). Across all 26,335 edges, T1w/T2w and *MBP* similarity were significantly correlated (Spearman r = 0.37), and correlations were lowest for regions highest in the cortical hierarchy (Spearman r = - 0.46, p_variogram_ < 0.001, **Figure 2E**).

Studies using bulk transcriptomic data on human cortex have found that: (1) mean T1w/T2w correlates strongly with the first principal component of gene expression (Burt et al. 2018); and (2) macroscale structural similarity is correlated with gene co-expression (Sebenius et al. 2023). We therefore hypothesised that T1w/T2w MIND similarity would be correlated with gene co-expression. To assess this hypothesis, we used single nucleus RNA sequencing data (Krienen et al. 2023) from 12 cortical areas (**Figure 3A**, **Supplementary Table 3**). The use of single cell transcriptomic data allowed us to assess the relationship between T1w/T2w MIND and gene co-expression within distinct cell types while avoiding cell type compositional differences as a potential confounder (Repsilber et al. 2010). We estimated whole genome co-expression matrices for 6 cell types (**Figure 3A**) and correlated these with a matched average adult T1w/T2w MIND network (N=340, mean age = 4.3 years). The cell type-specific correlation between whole genome co-expression and MIND similarity was strongest for glutamatergic neurons (Spearman r = 0.43; uncorrected p = 0.038); although no cell types remained significant after correction for multiple (6) comparisons (**Figure 3B**). When we repeated this analysis substituting *MBP* MIND edge weights for T1w/T2w MIND edge weights, we found much stronger relationships with cell-type specific genome co-expression. Inter-areal co-expression by glutamatergic neurons was most strongly correlated with *MBP* MIND (Spearman r = 0.67, p_FDR_ < 0.001). Whole genome co-expression was significantly correlated with *MBP* MIND in all neuronal and macroglial cell types after correction for multiple comparisons (**Figure 3C**).

**Figure 3:**
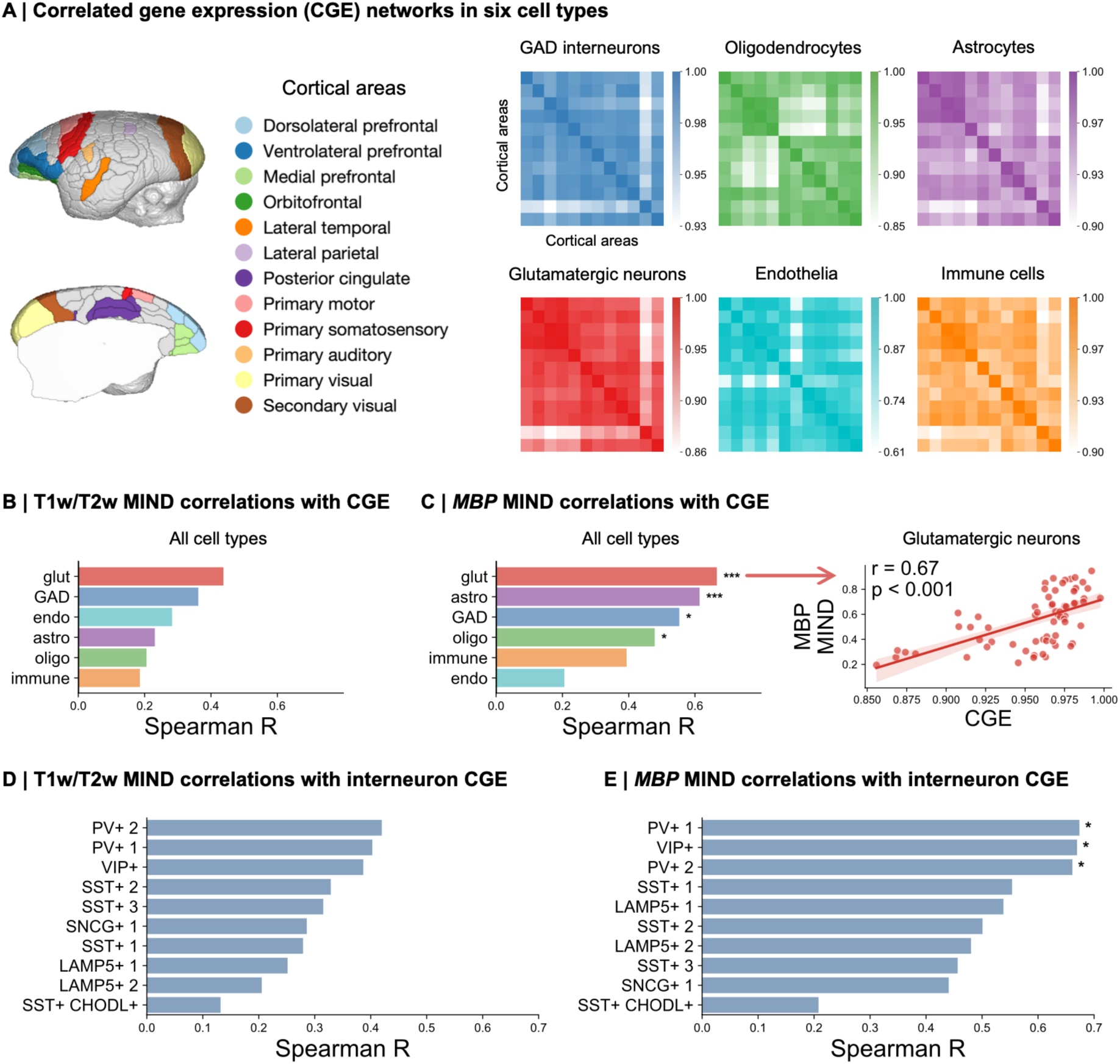
Myeloarchitectonic similarity is associated with co-expression of genes in glutamatergic neurons and PV+ and VIP+ interneurons. **A** Cell type- specific gene co-expression matrices estimated using single nucleus RNA sequencing data from 12 cortical areas (Krienen et al. 2023). In the following plots, bars represent the correlation (Spearman’s rho) between myeloarchitectonic (T1w/T2w or MBP expression) similarity networks and whole genome co-expression networks. Asterisks indicate the level of significance after comparison with spatial autocorrelation preserving nulls and Bonferroni correction for multiple comparisons: * p < 0.05, ** p < 0.01, *** p < 0.001. **B** T1w/T2w MIND is most strongly associated with whole genome co-expression by glutamatergic neurons. **C** MBP MIND was significantly correlated with whole genome co-expression by all neuronal and macroglial cell types, most strongly with glutamatergic neurons. **D** Both T1w/T2w MIND and MBP MIND were most strongly correlated with gene co-expression in PV+ and VIP+ interneurons, with MBP correlations remaining significant after correction for multiple comparisons.

Cortical cell types contain considerable heterogeneity (Krienen et al. 2020, 2023). We therefore hypothesised that there may be cell subtypes driving the relationship between gene co-expression and MIND similarity. We elected to test this relationship within GAD+ interneurons as previous work has indicated that markers of distinct interneuron subtypes are differentially correlated with T1w/T2w and other measures of cortical hierarchy in diverse species, with parvalbumin (PV+) interneurons being particularly strongly co-located with cortical hierarchy (Kim et al. 2017; Burt et al. 2018; Fulcher et al. 2019; Chen et al. 2023). We therefore estimated the correlations between interneuronal subtype-specific whole genome co-expression and T1w/T2w MIND edge weights (as in **Figure 3B-C**) for a set of 10 interneuron subtypes defined a priori by hierarchical clustering (Krienen et al. 2023, **Supplementary Figure 8**). We observed that both T1w/T2w MIND and *MBP* MIND were most strongly correlated with whole genome co-expression by PV+ and vasoactive intestinal peptide (VIP+) interneuronal subtypes (**Figure 3D**). Correlations between T1w/T2w MIND and genome co-expression by type 2 PV+ (Spearman r = 0.42, uncorrected p_variogram_ = 0.034) and VIP+ (Spearman r = 0.37, uncorrected p_variogram_ = 0.047) interneurons were nominally significant. Correlations between *MBP* MIND similarity and whole genome co-expression by PV+ and VIP+ interneurons remained significant after correction for multiple comparisons.

### Age-related changes in cortical T1w/T2w

We modelled age-related changes in a subset of N=210 marmosets with ages ranging from 7 to 36 months (pre-puberty to mature adulthood; **Figure 4A**). We used multiple linear regressions to estimate the linear slope of mean T1w/T2w change over this age range (**Figure 4B**). Corroborating previous findings in humans (Baum et al. 2022; Boroshok et al. 2023), we found that primary sensory regions lower in the cortical hierarchy showed greater age-related increases in T1w/T2w, while regions higher in the hierarchy showed smaller age-related increases in T1w/T2w (Spearman r = -0.61, p_variogram_ = 0.05, **Figure 4C**), though this trend did not reach significance. Furthermore, β_age_ coefficients were significantly positively correlated with pre-pubertal (< 1 year) mean T1w/T2w (Spearman r = 0.69, p_variogram_ < 0.001), indicating that regions which are more highly myelinated before puberty become relatively more myelinated over the course of adolescence: a “rich get richer” pattern of myelination (**Supplementary Figure 9A**). β_age_ coefficients were robust to the inclusion of covariates acting as additional T1w/T2w image normalisations (**Supplementary Methods**, **Supplementary Figure 9B**). Furthermore, these results were not unique to MRI; rate of change in mean T1w/T2w was highly correlated with the difference in mean MBP expression between a 6-month-old (pre-pubertal) male and 4-year-old (mature adult) male animal (Spearman r = 0.60, p_variogram_ < 0.001, **Figure 4D**).

**Figure 4:**
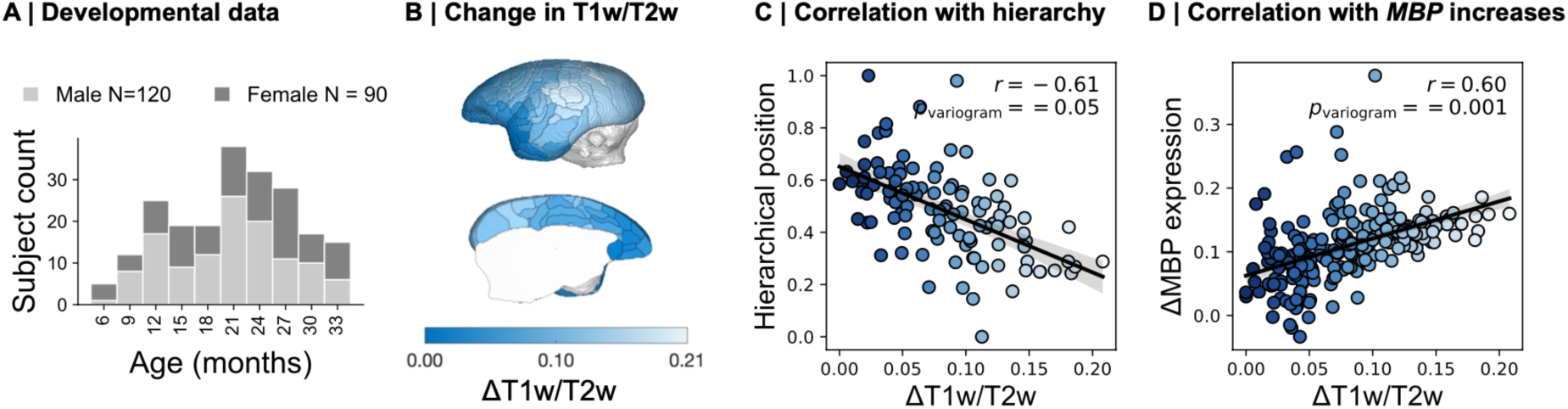
Marmosets show conserved adolescent increases in myelination. **A** Animals used for developmental analysis ranged from 7 months (pre-puberty) to 3 years (mature adulthood). **B** Rate of change in mean T1w/T2w for each region during this period, assessed using multiple linear regressions. **C** Rate of change in mean T1w/T2w is negatively associated with a region’s position in the cortical hierarchy. **D** Rate of change in mean T1w/T2w is significantly correlated with change in mean MBP expression.

### Developmental changes in myelination similarity

We hypothesised that T1w/T2w MIND would also show age-related changes representative of the inter-areally coordinated maturation of cortical myeloarchitecture. To qualitatively assess whether T1w/T2w MIND networks showed age-related changes, we assigned individual networks for animals aged <3 years to one of 7 age-defined bins (each spanning ≥4 months) and estimated the mean network for each age bin and the correlations between all pairs of averaged networks (**Figure 5A**). Correlations between networks progressively decreased as the separation in age between them increased, suggesting that there were indeed graded network changes occurring during development. After 20 months, roughly corresponding to the termination of adolescence (Sawiak et al. 2018), correlations between networks were more robust to variation in the age difference between them indicating a slowing or termination of developmental change in T1w/T2w MIND. This maturational plateau was consistently found to begin between 18-21 months in sensitivity analyses using bin sizes of 3 and 6 months (**Supplementary Figure 10**).

**Figure 5:**
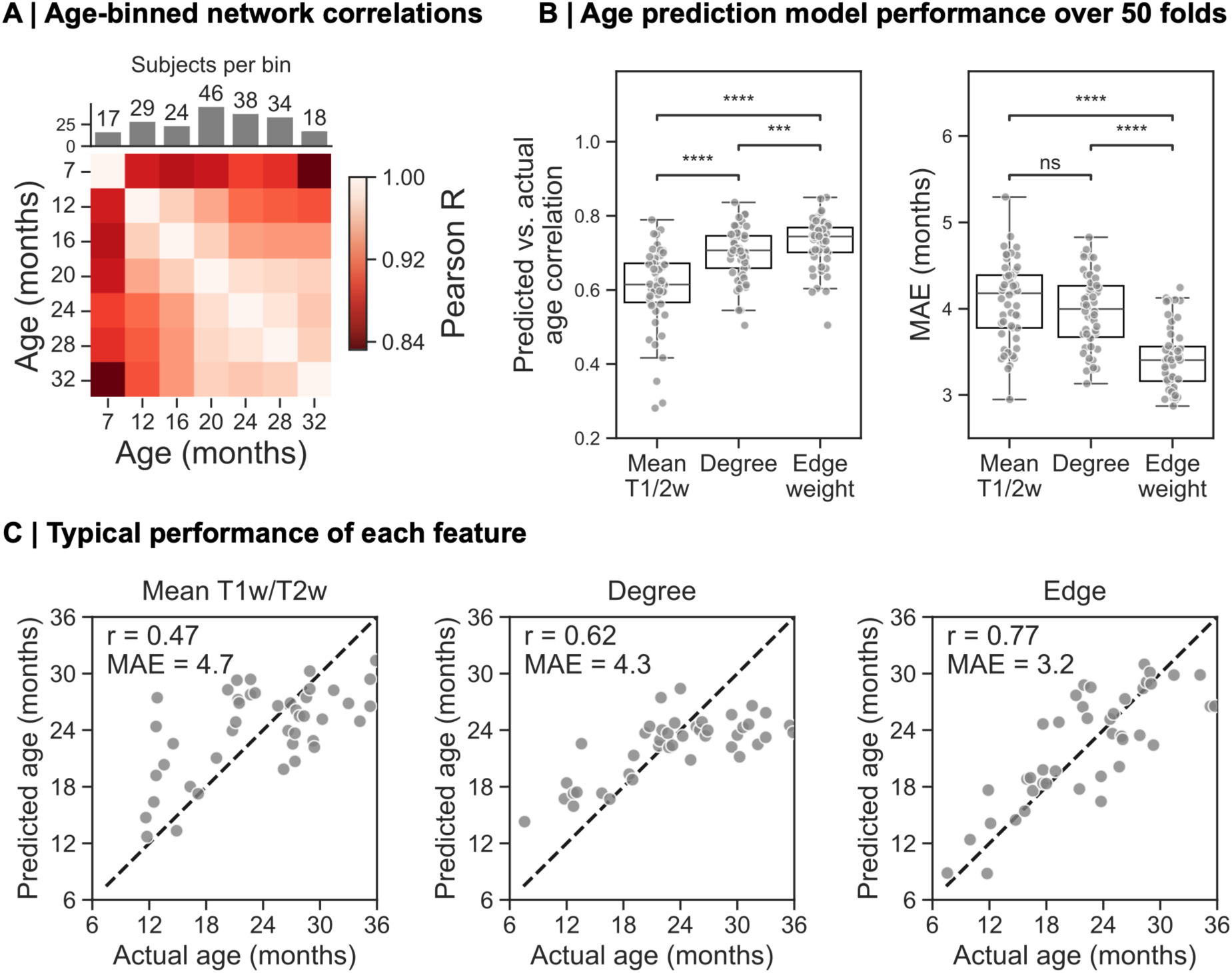
T1w/T2w MIND network edges and nodal degrees predict age more accurately than regional mean T1w/T2w. **A** Networks averaged into ≥4-month age bins become decreasingly correlated as the age between them increases. Correlations remain high after 20 months, indicating that there is relatively little change in T1w/T2w similarity beyond this point. **B** Performance of age prediction models trained on mean T1w/T2w and network features (degree and edge weights) over 50 splits of the data. **C** Typical model performance for each predictor. Performance of the model with the median partial R across 50 folds is shown here.

To assess the extent to which T1w/T2w MIND networks capture developmental changes which are chronologically consistent across individuals, we used machine learning models to predict subject age. Models were trained to predict age from an individual scan using three distinct sets of features: 1) regional mean T1w/T2w signal itself, 2) MIND networks summarised by the weighted degree of each node and 3) all edge weights in the MIND network. We measured performance using the partial correlation between predicted and true age, corrected for sex and eTIV (Sebenius et al. 2023), and the mean absolute error (MAE) between predicted and true ages. Over 50 splits of the data, T1w/T2w MIND network features out-performed regional mean T1w/T2w in predicting age (**Figure 5B**), indicated by higher partial correlations and lower MAE. Example model predictions (**Figure 5C**) illustrate a plateau in age predictions around 2 years, evident in the degree-based prediction. MIND edge weights and nodal degree gave more accurate predictions than regional mean T1w/T2w, supporting the added value of MIND network metrics in characterising intracortical maturation of myelin.

To identify specific age-related changes in nodal degree and edge weights of T1w/T2w MIND networks that were driving the superior predictive performance of MIND network features (**Figure 5**) we used multiple linear regression. At the degree level, 73/230 regions showed significant linear changes after correction for multiple comparisons (**Supplementary Figure 11A**). Anterior temporal, mainly auditory, areas showed significant degree decreases, reflecting an increasingly differentiated myeloarchitectonic structure relative to the rest of the cortex (**Supplementary Figure 11B**). Conversely, some somatosensory, primary motor, and visual areas showed significant degree increases, reflecting myeloarchitectonic convergence with the rest of the cortex (**Supplementary Figure 11B**). This pattern of changes was robust to the addition of estimated total intracranial volume as a covariate (**Supplementary Figure 11C**).

We performed the same regression analyses at each edge. The β_age_ coefficient of each edge, representing the linear rate of change in similarity between each pair of cortical nodes, is shown in **Figure 6A**, grouped into cortical zones. This revealed bidirectional changes in similarity of certain cortical areas which were not apparent in the degree-level analyses. For example, orbitofrontal cortex (OFC) showed decreases in similarity to posterior parietal (PPC), auditory (AUD), and visual (VIS) cortices, and increases in similarity to dorsolateral prefrontal (DLP) and anterior cingulate (ACC) cortices. Mean age-related changes between pairs of cortical zones is visualised in **Supplementary Figure 12A**.

**Figure 6:**
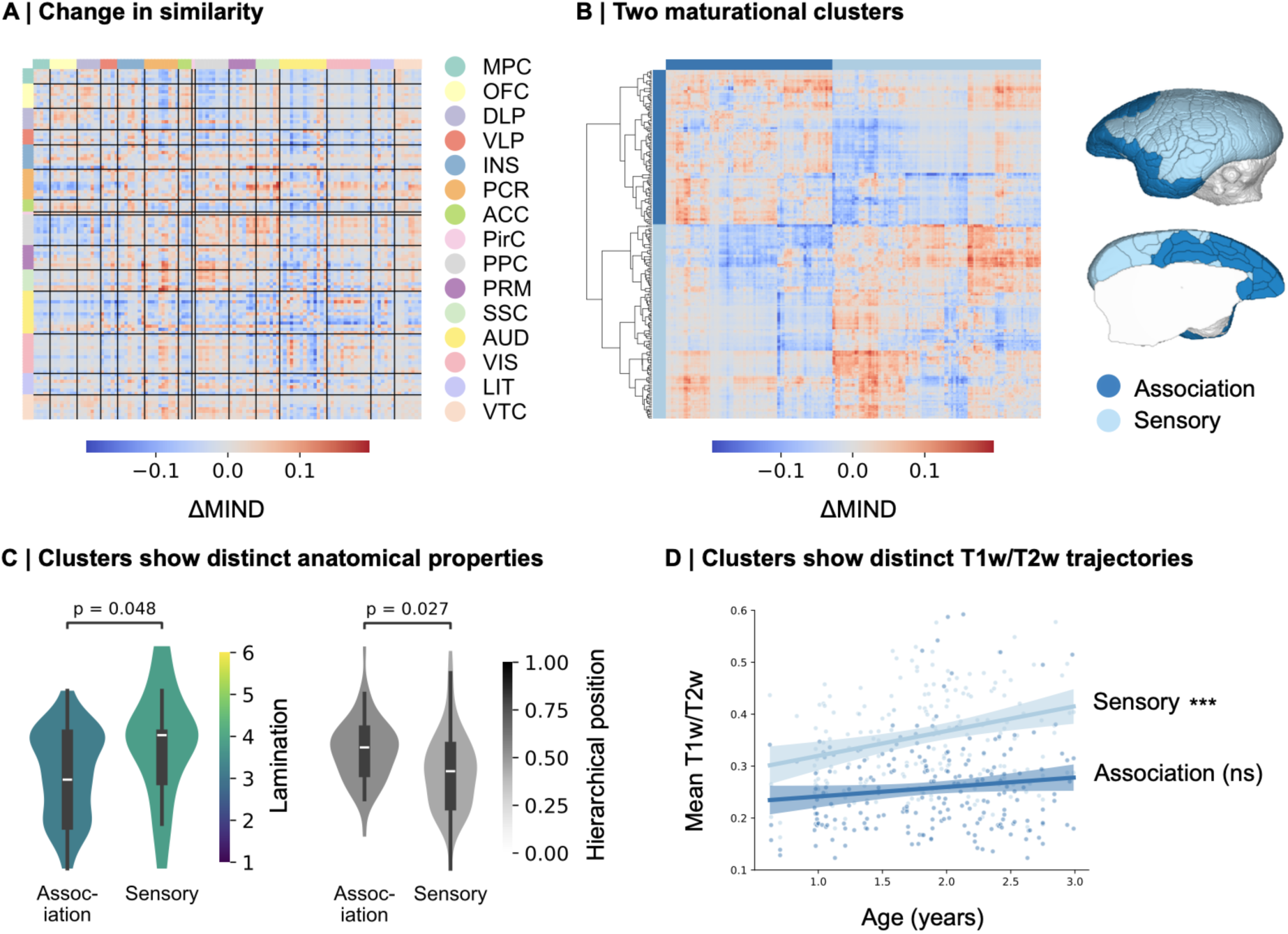
Sensory and association cortex show distinct age-related changes in myelination network properties. **A** Annual change in T1w/T2w similarity between all pairs of cortical areas grouped by cortical zones. Matrix shows the left hemisphere. **B** Hierarchical clustering reveals two cortical clusters with similar patterns of maturation. Matrix shows both hemispheres. **C** The sensory cluster had significantly higher mean degree of lamination and significantly lower mean hierarchical position. Reported p- values were derived from two-tailed, two-sample t-tests evaluated against spatially autocorrelated null models. **D** Maturational clusters were differentiated by their linear mean T1w/T2w trajectories. Asterisks represent the statistical significance of the β_age_ coefficient in multiple linear regressions predicting mean T1w/T2w of each cluster using age and sex as covariates.

We next used agglomerative hierarchical clustering as a data driven approach to explore how cortical areas were coordinated in terms of their adolescent maturation. This approach iteratively groups cortical areas into clusters based on how statistically similar their pattern of edge-level changes are. The two-cluster solution separated the cortex into one cluster containing mainly primary sensory and motor areas (“sensory”) and one cluster containing mainly frontal and paralimbic association areas (“association”: **Figure 6B, Supplementary Figure 12B**). In keeping with these identities, clusters were anatomically differentiable, with sensory cluster having significantly higher mean degree of lamination (t = 2.46, p_variogram_ = 0.048) and significantly lower mean hierarchical position (t = -2.98, p_variogram_ = 0.027; **Figure 6C**). Clusters also showed distinct mean T1w/T2w trajectories, the sensory cluster becoming significantly more myelinated during adolescence and increasingly divergent from the association cluster (**Figure 6D**). When repeating this analysis using three or four clusters, the association cluster remained largely intact, suggesting that its constituent regions are strongly maturationally coupled (**Supplementary Figure 12C**).

We replicated edge-level findings using *MBP* expression data. Here, developmental change was approximated using the difference between a mature adult (4-year-old) and pre-pubertal (6-month-old) male *MBP* MIND network (**Supplementary Figure 13A**). This revealed large decreases in similarity between primary sensory and prefrontal cortical regions (**Supplementary Figure 13B**). Clustering of the *MBP* MIND developmental matrix aligned with the T1w/T2w MIND developmental matrix, yet also reflected modality-specific developmental trends. This was evidenced by a moderate cophenetic distance correlation (Spearman r = 0.20; **Supplementary Methods**) and differences in overlap of cortical zones with clusters (**Supplementary Figure 13C-D**). However, *MBP* MIND developmental clusters demonstrated the same pattern of anatomical differences between clusters (**Supplementary figure 15E**), supporting overall correspondence between the two modalities in estimating developmental changes in cortical myelination networks.

## Discussion

We estimated structural similarity networks from myelin-sensitive MRI images in the common marmoset and largely confirmed our prior hypotheses: (1) that MRI-derived cortical similarity networks are closely co-located with commensurate maps derived from microscopic measurements of myelin basic protein gene expression and transcription at the cellular level; and (2) that MRI-derived cortical myeloarchitectonic similarity shows developmental changes which are stable across individuals and are different for regions occupying different levels of the sensory-association hierarchy, as in humans. We discuss these hypotheses in light of the existing literature.

T1w/T2w ratio calculation enhances contrast related to myelin in humans (Glasser and Van Essen 2011). We show, via co-location with spatially resolved brain images of myelin basic protein mRNA, that this applies to T1w/T2w images obtained from common marmosets at high magnetic field strengths. T1w/T2w and *MBP* expression similarity network edges were highly correlated, most so for edges to primary sensory and motor areas lowest in the cortical hierarchy. The reduced cross-modal concordance of similarity in higher-order cortical areas may arise due to their prolonged developmental and enhanced adult plasticity (Glasser et al. 2014; Sydnor et al. 2021), which would render their myeloarchitectonic structures more sensitive to individual-specific environmental influences and thus increase inter-individual variability in similarity of these areas.

We found that cortical areas with high myeloarchitectonic similarity also had similar whole genome expression profiles in neurons. We interpret this finding in two ways. First, axonal connectivity is guided by the degree of matching between specific patterns of adhesion molecules on neuronal cell surfaces (Sperry 1963; de Wit and Ghosh 2016) and axonally connected neurons typically exhibit greater similarity in their whole genome expression profiles (Fornito, Arnatkevičiūtė, and Fulcher 2019). Thus, if we interpret similarity as a weak proxy of axonal connectivity emerging due to homophily (Sebenius, Dorfschmidt et al. 2025) – supported by a moderate (r = 0.34) correlation with tract tracing – similarity and neuronal gene co-expression should also be correlated. Second, despite oligodendrocytes being the key cell type responsible for myelination in the brain, myelination is also dependent on neuronal factors, including the expression of permissive surface molecules and neuronal activity (Emery 2010). This interdependence may establish a coupling between local myeloarchitecture and neuronal gene expression patterns.

The coupling between myeloarchitectonic similarity and neuronal gene co-expression was strongest for glutamatergic and PV+ and VIP+ interneurons. Previous work has shown that transcriptomic markers of glutamatergic neurons and interneurons (most strongly PV+ but also VIP+ and SST+) correlate strongly with mean T1w/T2w (Burt et al. 2018; Fulcher et al. 2019) and other indices of cortical hierarchy (Kim et al. 2017; Chen et al. 2023). These neuronal subtypes form tightly anatomically and functionally interconnected cortical microcircuits (Douglas, Martin, and Whitteridge 1989; Pfeffer et al. 2013) that strongly reflect local laminar structure (Beul and Hilgetag 2014). Cortical areas with more similar myeloarchitecture are therefore likely to contain phenotypically similar cortical microcircuits and similar gene expression profiles in the cell types forming these circuits.

Our second main hypothesis centred around adolescence, the age range with peak onset of psychiatric disorders (Solmi et al. 2022) and during which increases in cortical myelin contribute to the closure of critical windows of enhanced sensitivity – and vulnerability – to environmental stimuli (Larsen et al. 2023). Previous studies have shown that linear increases in mean T1w/T2w vary along the cortical hierarchy, with primary sensory areas showing steepest increases and association areas showing shallowest increases over the course of adolescence (Baum et al. 2022; Boroshok et al. 2023). For the first time, to our knowledge, we show that this pattern is conserved in the common marmoset, highlighting its translational utility as a primate neurodevelopmental model.

We further demonstrate that T1w/T2w ratio similarity networks show adolescent changes which are stable across individuals and can therefore predict age. Previous work has shown that MIND networks generated from multiple microscale and macroscale MRI features can predict adolescent age in humans (Sebenius et al. 2023). Our work generalises this finding to a simpler context, using a single clinically ubiquitous MRI feature in a primate model. We found that network features predicted age more accurately than mean T1w/T2w alone, suggesting that relative myeloarchitectonic changes between cortical areas are more developmentally informative than absolute changes in mean myelin. Though the enhanced performance of edge-based prediction is unsurprising due to the large number of edges, it is striking that degree-based predictions, with one value per cortical area, out-perform mean T1w/T2w. A prior study using morphometric similarity networks (Seidlitz et al. 2018) estimated by pairwise correlations of 7 mean macrostructural features found that network features were *poorer* at predicting human adolescent age than mean cortical thickness or volume alone (Griffiths-King, Wood, and Novak 2023). Though our study did not use the same features, our results suggest that when using voxel-level information to estimate similarity, networks contain enhanced developmental sensitivity relative to regional mean values.

Similarity network changes were organised along an axis from primary sensory to transmodal association areas (Mesulam 1998; Sydnor et al. 2021). Cortical areas occupying similar positions along this axis showed similar patterns of edge level maturation, with particularly strong maturational coupling observed within a rostromedial transmodal association cluster that overlapped a putative human default mode network homologue identified by previous analysis of tract tracing data (Buckner and Margulies 2019). In human cortex, myelination networks estimated as the pairwise correlations between regional MTsat depth profiles showed conserved adolescent reorganisation, with developmental changes organised along a latent axis mirroring the sensory-association gradient (Paquola et al. 2019). MTsat profiles of primary sensory (idiotypic) cortex became increasingly divergent from paralimbic cortex, and intermediate areas became either more idiotypic-like or paralimbic-like, indicating a reinforcement of the sensory-association axis. Consistent with this, we observed that areas occupying distant locations in sensory-association space tended to diverge in their myeloarchitecture, with large decreases in similarity between primary sensory areas and prefrontal, temporal, and paralimbic association areas. Parallel evidence from rat cortex indicates that prefrontal areas become myeloarchitectonically differentiated from primary sensory areas between late adolescence and mid adulthood in MIND similarity networks estimated from myelin-sensitive Magnetisation Transfer Ratio MRI images (Smith et al. 2024). Taken together, these findings may indicate that the sensory-association axis represents a conserved gradient of myelination network reorganisation during adolescence, with areas co-localised along this gradient a) showing similar patterns of change to all other cortical areas and b) becoming themselves convergent in terms of their myeloarchitecture.

Our study contains limitations. Although sensitive to myelination, T1w/T2w ratio is not a direct measure of myelin. Hence, though we observed strong concordance between mean T1w/T2w and mean *MBP* expression across the cortical surface, there were differences in the signal distribution shapes (standard deviation and skewness) between MRI and *MBP* histology. These differences may arise because: a) though T1w/T2w ratio calculation enhances contrast related to myelin, images are sensitive to non-myelin factors such as iron and water content (Glasser and Van Essen 2011; Uddin et al. 2019) and b) noise and partial voluming effects introduced during MRI acquisition and preprocessing may influence regional T1w/T2w distribution shapes, particularly at the white matter and pial surfaces. These factors may contribute to the differences we observed in myeloarchitectonic network structure when estimated using MRI or *MBP* expression, which in turn may explain why we observed only nominally significant correlations between T1w/T2w similarity and gene co-expression. Here, we further anticipate that expanding the number of cortical areas beyond 12 (and the number of edges beyond 66) would allow for a finer-grained comparison between structural similarity and gene co-expression.

Second, we used linear modelling approaches as an interpretable way to model age-related changes, though we expect that cortical myelination similarity would show non-linear trajectories. In humans, developmental increases in cortical mean T1w/T2w follow non-linear trajectories, with primary sensory areas reaching peak myelination first, and association areas last (Grydeland et al. 2019; Baum et al. 2022). A comparable chronology has been observed for grey matter volume decreases in marmoset cortex (Sawiak et al. 2018), thought to reflect progressive increases in cortical myelination (Natu et al. 2019). Furthermore, cortical myelin undergoes dynamic changes across the lifespan. Indeed, myelination begins *in utero* for sensory and motor areas (Flechsig 1901, Yakovlev and Lecours 1967), while ageing involves cortical *de*-myelination. The regional sequencing of ageing-related myelin decreases is thought to be the chronological reverse of the sequence of adolescent myelin increases, reflecting a “first in last out” hypothesis of senescence (Grydeland et al. 2019). The field would benefit from an exploration of non-linear lifespan trajectories of myelination network features, for example using normative modelling approaches (Bethlehem et al. 2022), and from an investigation of how these trajectories relate to brain function, cognition, and psychiatric outcomes.

## Methods

### T1w/T2w ratio data and preprocessing

We used T1-weighted (T1w) and T2-weighted (T2w) images acquired at 9.4T from the Brain/MINDS project, currently the world’s largest open-access marmoset MRI data resource (Hata et al. 2023). The dataset contains cross-sectional MRI images from N=446 marmosets, spanning 0.62 to 12.75 years (N female = 238). Volumetric images were parcellated according to the 2019 Brain/MINDS Atlas (BMA 2019; Woodward et al. 2018), which is based on the histological atlas defined by Paxinos et al. (2012). Cortical areas used in our analyses are listed in **Supplementary Table 1** and the full preprocessing pipeline is visualised in **Supplementary Figure 1**. Briefly, T2w images were skull stripped using BrainSuite18a (Shattuck and Leahy 2002) to improve the quality of subsequent image registration, for which we used advanced normalisation tools (ANTs: Avants et al. 2011). An ex-vivo T2w template (Woodward et al. 2018) was warped to each T2w image, after which the warp was applied to the BMA 2019 atlas to generate a label image for each subject. Voxel-wise division of the T1w image by the T2w image generated a T1w/T2w ratio image which increased myelin-related contrast and reduced intensity inhomogeneity (bias; Glasser and Van Essen 2011). T1w and T2w images were not co-registered prior to calculation of T1w/T2w images because marmosets were anaesthetised and head-fixed before scanning, minimising head motion between T1w and T2w image acquisitions.

### Similarity network estimation using Morphometric Inverse Divergence

For each parcellated T1w/T2w image, a distribution of voxel values was extracted from each cortical area. Similarity between each pair of voxel value distributions was estimated using Morphometric Inverse Divergence (MIND; Sebenius et al. 2023) yielding a univariate (single feature) similarity network. MIND is the inverse of the Kullback-Leibler (KL) divergence, a statistical measure of divergence or dissimilarity, between two distributions (Kullback and Leibler 1951). For two sets of voxels (or vertices from a surface reconstruction) in cortical areas *a* and *b* following distributions *P_a_* and *P_b_*, the KL divergence between them is calculated as:

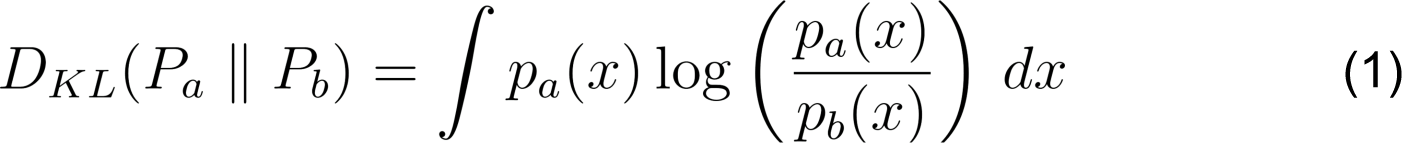

Where *p_a_(x)* and *p_b_(x)* are the probability densities at particular values of x. KL is then normalised and inverted, converting it to a measure of similarity:

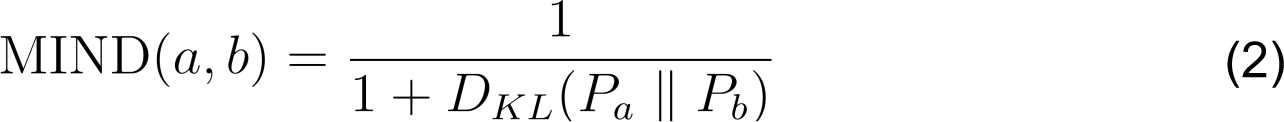

Estimation of MIND for each possible pair of areas in the cortical parcellation results in a similarity matrix, where each element (edge) represents the similarity in voxel distributions between a pair of cortical areas, ranging from >0 to 1. Low values indicate a highly dissimilar, divergent or differentiated pair of cortical areas and high values indicate a highly similar, convergent, or near-identical pair of cortical areas.

Two elementary features of each MIND network were the focus of further analysis: nodes and edges. Edge weights represent the pair-wise similarity between cortical nodes, and the weighted degree of each node is simply the average of the weights of all edges linking that node to the rest of the cortical network.

### Estimation of KL divergence

Non-parametric estimation of the KL divergence between two (voxel) distributions requires a systematic comparison of probability densities, which can be done either globally or locally. Global methods involve comparing coarse partitions of the distributions, for example corresponding histogram bins, while local methods involve comparing densities in the local neighbourhoods surrounding individual distribution values, i.e. voxel values (**Supplementary Methods**). Though both global and local KL estimation methods have previously been used to estimate structural similarity networks from MRI data (Kong et al. 2014; Sebenius et al. 2023), they have not been directly compared. We addressed this gap in the literature by systematically comparing a global and local KL estimator. We used a global estimator of KL which binned the data into successive histograms (Wang, Kulkarni, and Verdu 2005) using in-house code. Secondly, we used the local estimator implemented in the current MIND algorithm (Sebenius et al. 2023) based on a k-Nearest Neighbours (k-NN) approximation (**Supplementary Methods**).

To facilitate comparisons between similarity networks generated using different KL estimators, and variable parameters of each of those estimators, we adopted a previously used set of anatomical benchmarks (Sebenius et al. 2023). Anatomical benchmarking assumes that the “best-performing” or “most optimal” MRI similarity networks are ones that correspond most strongly with the underlying anatomy. In our case, benchmarks were based on cytoarchitecture, interhemispheric symmetry, and axonal connectivity (**Supplementary Methods**). We additionally compared MRI similarity networks resulting from these KL estimation algorithms using an age prediction task. Here, the networks producing most accurate age predictions, assessed via their correlation and mean squared error relative to actual ages, were deemed “most optimal”.

### Marmoset anatomical data

#### Cytoarchitecture

We used the cytoarchitectonic class assignment map of Atapour et al. (2024), in which cortical regions were assigned to one of six classes based on expert classification of histological sections (**Supplementary Figure 5B**). In order of increasing lamination, the classes are: Agranular (Ag), Dysgranular (Dys), Eulaminate I (Eu I), Eulaminate II (Eu II), Eulaminate (Eu III), and Koniocortex (Kon).

#### Axonal connectivity

Tract tracing data were obtained from Majka et al. (2020). Data from 143 retrograde tracer injections in 52 adult marmosets (N female = 21, aged 1.4-4.6 years) were used to generate a single {55 x 116} tract tracing matrix. Retrograde tracer injections were placed in 55 target regions, labelling axonal tracts originating from cell bodies in 116 source regions. Quantitative analysis of histological sections was used to estimate the fraction of labelled neurons (FLN) originating in each source brain region for each of the 55 target regions receiving a tracer injection. The logarithm of the estimated fraction of labelled neurons in each region, Log_10_(FLN), averaged across animals, was used as the key metric of axonal connectivity. Log_10_(FLN) ranges from 0 (100% of tracts originate in source) to -∞ (0% of tracts originate in source). Because this approach fails to capture axonal pathways terminating in the 61 regions not injected with tracer, we used the edge-complete matrix of {55 x 55} regions that had received at least one injection. For comparability, MIND similarity matrices were restricted to the same 55 regions in all analyses involving tract tracing data.

#### Position in the cortical hierarchy

A cortical area can be assigned to a position in the cortical processing hierarchy based on its laminar pattern of incoming and outgoing axonal connections (Felleman and Van Essen 1991). This is based on the observation that feedforward connections, carrying information up the cortical hierarchy from sensory to association regions, arise mainly from supragranular layers and terminate in layer IV (Rockland and Pandya 1979). The proportion of tracts a cortical area receives which originate in supragranular layers has therefore been used to estimate cortical maps of hierarchical position in the marmoset (Theodoni et al. 2021), with regions receiving a high proportion of feedforward information being located high in the hierarchy and regions receiving a low proportion of feedforward information being located low. We used the hierarchical position map from Theodoni et al. (2021), which assigns to 110 of 115 cortical areas a value ranging from 0 (bottom of the hierarchy: primary visual cortex) to 1 (top of the hierarchy: agranular insula). This map is visualised in **Supplementary Figure 5C**.

### Marmoset transcriptional data and pre-processing

#### In-situ hybridisation data

We downloaded spatially resolved images of myelin basic protein (*MBP*) messenger RNA (mRNA) from the Marmoset Gene Atlas (Shimogori et al. 2018; Kita et al. 2021). Images were collected using in-situ hybridisation (ISH), a histochemical technique allowing spatial localisation of specific nucleic acids within a tissue sample (Wilcox 1993). We downloaded in-situ hybridisation images of *MBP* mRNA from a 6-month-old (pre-pubertal) male and a 4-year-old (mature adult) male marmoset. For both animals, *MBP* mRNA was measured in 90 coronal brain slices, spaced 196 μm apart (Shimogori et al. 2018; Kita et al. 2021). Though the posterior pole of occipital cortex and the anterior pole of frontal cortex were not completely covered, each brain region in the Paxinos et al. (2012) parcellation had at least one representative tissue sample analysed at both ages.

To facilitate a quantitative comparison of T1w/T2w and *MBP* mRNA images, in-situ hybridisation slices were stacked to form a volume, which was then linearly and nonlinearly deformed to the BMA 2019 T2w template. To do this, we implemented a preprocessing pipeline, fully described in **Supplementary Figure 6**, that used a combination of in-house code and code made available from Tong et al. (2022). This pipeline allowed us to estimate the distribution of *MBP* mRNA expression in each cortical area, with comparable spatial resolution to the T1w/T2w images (isotropic voxel resolution of *MBP* mRNA images = 0.10mm; T1w/T2w images = 0.27mm). The resulting cortical maps were subsequently used to estimate MIND networks representing inter-areal similarity of *MBP* expression on the same spatial scale and using the same areal parcellation as the MIND networks derived from T1w/T2w images.

#### Single-cell, whole genome mRNA data

We used open-access single nucleus RNA sequencing data from Krienen et al. (2023). This dataset contains whole genome transcripts from multiple cortical and subcortical regions comprising six cell types: glutamatergic excitatory neurons (glut), glutamic acid decarboxylase-expressing inhibitory interneurons (GAD), oligodendrocytes (oligo), astrocytes (astro), microglia and macrophages (immune), and endothelial cells (endo). Further details on data collection, quality control, and cell type classification are provided by Krienen et al. (2023).

Cortical dissections in Krienen et al. (2023) were guided by the Paxinos et al. (2012) atlas. Some dissections were restricted to individual Paxinos regions, while others were inclusive of several neighbouring regions, constituting a larger amalgamated area (Krienen, personal communication). This resulted in a partial parcellation of cortex into 12 relatively coarse-grained areas (**Figure 3A**, **Supplementary Table 3**). We used this scheme to coarse-grain the more finely parcellated T1w/T2w and *MBP* mRNA images so that they were anatomically aligned with the cortical regions for which single-cell sequencing data were available. Further details on the coarse parcellation can be found in **Supplementary Table 3**, and the sex and age of the sampled animals are described in the figure legend.

We used these data to generate cell type-specific correlated gene expression (CGE) matrices. Cell-by-gene matrices for each cell type were downloaded from the CELLxGENE data repository (RRID:SCR_021059). Library size normalisation, i.e. normalising gene expression across cell types to correct for differences in sequencing depth, had already been done by Krienen et al. (2023). We retained the top 10% most variable genes for CGE network calculation using Scanpy’s highly_variable_genes function (Wolf, Angerer, and Theis 2018). Our reasoning was that genes with low variability in expression across the cortex are unlikely to drive systemic variation in cortical myeloarchitecture. Indeed, previous work has shown that transcriptomic variation in a subset of 200 genes that are most highly variable across cortex is closely coupled to variation in mean T1w/T2w of human visual cortex (Gomez, Zhen, and Weiner 2019). Within each cell type, we averaged log normalised expression across cells with the same regional annotation to form a region-by-gene expression matrix. Pearson correlations between each pair of regional gene expression vectors were used to generate the final CGE matrix for each cell type (**Figure 3A**). The same process was applied to interneuron subtypes to generate interneuron subtype-specific correlated gene expression matrices.

### Statistical analysis

#### Multiple linear regression

To estimate linear changes in nodal properties (mean T1w/T2w and degree), and edge weights during development, we used multiple linear regressions with age as the independent variable. Multiple comparisons testing was conducted using Benjamini Hochberg correction. We included sex as a covariate in all models as studies have demonstrated sex differences in regional cortical mean T1w/T2w in humans (Küchenhoff et al. 2024). Furthermore, some studies indicate that humans display sex-specific adolescent trajectories of cortical mean T1w/T2w (Norbom et al. 2020), therefore we tested for the significance of an age by sex interaction term in our models. However, this interaction term was ultimately dropped from all models as it did not reach significance in any model after multiple comparisons correction. Furthermore, sensitivity analyses revealed that addition of two normalisation covariates to our T1w/T2w models, and the addition of estimated total intracranial volume (eTIV), to degree models did not greatly change β_age_ values (**Supplementary Figure 9B, 11C**). For further discussion on covariate considerations, see **Supplementary Methods**.

#### Spatial null models

In spatially embedded systems, such as the brain, regions that are physically closer to each other tend to be more similar across a wide range of features, including axonal connectivity profiles, cytoarchitectonic structure, and gene expression (Fornito, Arnatkevičiūtė, and Fulcher 2019; Markello and Misic 2021). This spatial dependence, or autocorrelation, between neighbouring brain areas violates the assumption of independence in standard statistical inference procedures and can lead to inflated type I error rates if not properly addressed (Markello and Misic 2021). We therefore used spatial autocorrelation-preserving null models, generated using BrainSMASH (Burt et al. 2020), for statistical inference on parcellated cortical maps. BrainSMASH implements an algorithm that estimates for a particular brain map (for instance, mean T1w/T2w) the variance in regional values as a function of the Euclidean distance between regions (the variogram) and generates surrogate brain maps with matched variograms (Viladomat et al. 2014) that can be used to construct an appropriate null distribution of test statistics. BrainSMASH permuted maps were computed from left hemispheric data only. We used a similar procedure to assess the significance of correlations with similarity networks at the edge level (**Supplementary Methods**). All statistical tests used 1000 permutations.

### Age prediction

To assess the extent to which MIND networks capture developmental changes in myelination, we trained machine learning models to predict age from T1w/T2w scans in N=210 animals aged 7 months (pre-pubertal) to 36 months (mature adult). Repeated 5-fold cross validation with 50 random permutations of the data was used to compare age prediction performance for models trained using either i) mean T1w/T2w per cortical area, ii) MIND weighted degree per cortical area, or iii) MIND edges between each pair of cortical areas. Each fold was balanced for sex and subjects in the top or bottom 1% of mean T1w/T2w or MIND weighted degree (averaged over all cortical areas) were excluded to reduce the impact of outliers. For each outer test fold, models were evaluated using the mean absolute error (MAE) and the partial correlation between actual and predicted age values, corrected for sex and estimated total intracranial volume. For details on quality control, model specification, and model selection, please refer to **Supplementary Methods**.

## Data availability

All data analysed in this study are publicly available. Retrograde tract tracing data are available from Majka et al. (2020). Region-wise hierarchical position assignments are available from Theodoni et al. (2021). Region-wise cytoarchitectonic class assignments are available from Atapour et al. (2024). MRI data (Hata et al. 2023) are available from the Brain/MINDs 2.0 data portal (https://dataportal.brainminds.jp/marmoset-mri-na216). In-situ hybridisation data (Shimogori et al. 2018 and Kita et al. 2021) are available from the Brain/MINDs Marmoset Gene Atlas data portal (https://gene-atlas.brainminds.jp/). Processed single nucleus RNA-sequencing data (Krienen et al. 2023) are available from the CELLxGENE data repository (RRID:SCR_021059; https://cellxgene.cziscience.com/collections/0fd39ad7-5d2d-41c2-bda0-55bde614bdb). Marker gene sets used to assign biological identities to interneuron clusters are available from Bakken et al. (2021).

## Code availability

Preprocessing of MRI data used Brainsuite18a (Shattuck and Leahy 2002) and ANTs (Avants et al. 2011). Preprocessing of in-situ hybridisation data used a modified version of previously published code (Tong et al. 2022; https://github.com/TrangeTung/marmoset_gradient) and ANTs. Computation of MIND networks used previously published code (Sebenius et al. 2023; https://github.com/isebenius/MIND). Downstream analyses were conducted in python. Processing of single cell data used Scanpy (Wolf et al. 2018). All custom code used in this study will be made publicly available on GitHub upon publication of the final article.

## Acknowledgements

E.D.H was generously supported by a Rosetrees Trust grant MB2023\100002 and funding from the University of Cambridge School of Clinical Medicine. S.J.S. and A.C.R were supported by a Medical Research Council Programme grant MR/V033492/1. E.T.B. was supported by a National Institute for Health and Care Research (NIHR) Senior Investigator award.

## Author information

### Contributions

E.D.H. and E.T.B. conceived of the primary methodology. E.D.H. performed all analyses and drafted the manuscript. R.A.I.B., A.C.R., and E.T.B. provided supervision. S.J.S. advised on analyses of age-related changes. E.D.H., S.J.S., R.A.I.B., A.C.R., and E.T.B. reviewed and edited the manuscript.

