## Supplementary Information for "Cortical myelination networks reflect neuronal gene expression and track adolescent age in marmosets"

#### Contents


**Supplementary Methods** **2**

Local and global estimators of KL divergence 2
Edge-level effects of tuning parameters in KL divergence estimators 3
Anatomical benchmarks 3
Selecting optimal Histogram parameters using anatomical benchmarks 4
Statistical analysis 5
Age prediction 5
Comparing clustering solutions between modalities 6

**Supplementary Tables** **7**

1. Regions used in whole cortex analysis* 7
2. Tuneable parameters in KL estimators 8
3. Coarse-grained parcellation used in single cell transcriptomic analyses 9

* Table included as separate excel file

**Supplementary Figures** **10**

1. Preprocessing of T1w/T2w ratio images 10
2. Edge-level effects of tuning parameters in KL divergence estimators 11
3. Selecting optimal Histogram algorithm parameters 12
4. Local algorithm out-performs global on anatomical benchmarks and age prediction 13
5. Anatomical maps 14
6. Preprocessing of in-situ hybridisation images 15
7. Comparing T1w/T2w and MBP expression distribution shapes 16
8. Assignment of interneuron clusters to subtypes 17
9. Age-related changes in T1w/T2w 18
10. Adult maturational plateau is robust to bin size 19
11. Auditory regions become microstructurally differentiated from the rest of the cortex 20
12. Maturational clustering sensitivity analyses 21
13. Maturational changes in *MBP* expression similarity are concordant with changes in T1w/T2w ratio similarity 22

**Supplementary References** **23**

### Supplementary Methods

#### Local and global estimators of KL divergence

Non-parametric estimation of the KL divergence between two (voxel) distributions requires a systematic comparison of their probability densities (Noshad et al. 2017). In the univariate (single feature) case, this can be performed using histograms, kernel density estimation (KDE) or nearest neighbour methods (Silverman 2018). Histograms and KDE estimate a probability density function over all points in a distribution. We therefore refer to them as "global" estimators. For a pair cortical areas, a and b, following distributions $P_a$ and $P_b$, the probability densities at equivalent values of x are successively substituted into equation (1) to estimate KL:

$$D_{KL}(P_a \parallel P_b) = \int p_a(x) \log \left( \frac{p_a(x)}{p_b(x)} \right) dx \tag{1}$$

Where $p_a(x)$ and $p_b(x)$ are the probability densities at equivalent values of x, i.e. at equivalent bins if using histograms (Wang, Kulkarni, and Verdu 2005). Previous studies have used KDE to estimate KL divergence-based similarity networks from voxel-wise grey matter intensity (Kong et al. 2014, 2015) and grey matter volume (Wang et al. 2016). These studies have shown that univariate similarity networks exhibit high test-retest reliability (Wang et al. 2016) and are robust to variation of smoothing parameters and parcellation schemes (Wang et al. 2016). These networks also show bidirectional changes in grey matter similarity following sleep deprivation (Kong et al. 2014, 2015) and decreases in global efficiency and clustering of networks during healthy ageing (Kong et al. 2014, 2015).

Conversely, nearest neighbour methods involve comparisons of the local density around individual (voxel) values, rather than between relatively coarse partitions of the data (Silverman 2018). We therefore refer to them as "local" methods. MIND has previously been estimated with a k-Nearest Neighbours (k-NN)-based KL estimator developed by Perez-Cruz (2008). For two sets of voxels, $V_a$ and $V_b$, following distributions $P_a$ and $P_b$, KL is estimated as:

$$D_{KL}(P_a \parallel P_b) = -\frac{d}{n} \sum_{i=1}^{n} \log \frac{r_k(x_i)}{s_k(x_i)} + \log \frac{m}{n-1} \tag{2}$$

Where $d$ is the number of structural features measured at each voxel, $n$ and $m$ are the number of values (voxels) in $V_a$ and $V_b$ respectively, and $r_k(x_i)$ and $s_k(x_i)$ are the differences between the voxel value $x_i$ to the most similar voxel value in $V_a$ (its own distribution) and $V_b$ (the other distribution), respectively.

#### Edge-level effects of tuning parameters in KL divergence estimators

Histogram and k-NN estimators of KL divergence include several tuneable parameters. For the histogram algorithm, these are: the number of bins used to partition the data; epsilon, a small constant added to $p_b(x)$ in Equation (1) to prevent division-by-zero errors; and the percentiles of data retained when calculating histogram bins. For the k-NN algorithm, these are: *k*, the $k^{th}$ nearest neighbour used to estimate local density; and the percentiles of data retained. Full parameter descriptions and the default values used in our KL estimators are provided in **Supplementary Table 2**.

We reasoned that the choice of parameters may impact the estimated value of KL at a single edge and thus may impact the overall MIND network structure. We therefore undertook a series of simulations to assess which parameters had the largest impact on KL estimation across one edge and all network edges. At the single edge level, we estimated KL between two empirical T1w/T2w voxel distributions extracted from a single subject (**Supplementary Figure 2A**) using default parameter values. We then independently varied each KL estimation parameter in the k-NN and Histogram algorithm and calculated the percentage change in estimated KL from default (**Supplementary Figure 2B**). This revealed that the percentiles parameter for both algorithms and the number of bins used in the histogram algorithm had the largest effects on the value of KL. We then performed a similar analysis at the whole network level (i.e. over all edges), by first generating a full MIND network using default KL parameters (**Supplementary Figure 2C**) and assessing how modulating parameters altered the edge-wise correlations with the default network. This revealed, again, that the same parameters led to largest changes in network edge structure (**Supplementary Figure 2D**). Our simulations indicate that the percentiles parameter for both algorithms and the number of bins used in the histogram algorithm should be carefully selected prior to downstream analysis as they generate large changes in MIND network structure.

#### Anatomical benchmarks

To facilitate comparisons between similarity networks generated using different KL estimators, and variable parameters of each of those estimators, we adopted a previously used set of anatomical benchmarks (Sebenius et al. 2023). Anatomical benchmarking assumes that the “best-performing” or “most optimal” MRI similarity networks are ones that correspond most strongly with the underlying anatomy. In our case, benchmarks were based on cytoarchitecture, interhemispheric symmetry, and axonal connectivity.

*Cytoarchitecture*

One explanation for macroscale MRI similarity is a result of similarity at the microscale (Sebenius, Dorfschmidt et al. 2025). Therefore, we expect that regions of the same cytoarchitectonic class, due to having similar laminar organisation on microscopy, should tend to be similar in terms of their T1w/T2w voxel distributions. In this case, networks which have a higher proportion of intra-class edges over a range of densities, i.e. those with a higher area under the curve (AUC), are assigning higher macroscale T1w/T2w similarity to pairs of regions with higher microscale myeloarchitectonic similarity, which is favourable. We used the cytoarchitectonic class assignment map of Atapour et al. (2024), described in **Methods** and illustrated in **Supplementary Figure 5B**.

*Interhemispheric symmetry*

Though notable cortical asymmetry exists, homotopic regions are highly structurally similar due to the shared genetic effects of antero-posterior morphogen gradients during early patterning of the telencephalon (Sansom and Livesey 2009). Our second benchmark was therefore to assess the proportion of all (115) homotopic edges that were present across a range of network densities (Sebenius et al. 2023). Networks with a higher AUC are more optimally capturing high similarity between homotopic cortical areas.

*Axonal connectivity*

Due to the principle of homophily, regions with higher macroscale similarity are more strongly axonally connected (Sebenius, Dorfschmidt et al. 2025). We therefore expected biologically optimal networks to correlate with axonal connectivity over a range of network densities, and networks with a higher AUC are more optimally capturing homophilic attachment principles.

#### Selecting optimal Histogram parameters using anatomical benchmarks

Changing the parameters of KL estimation can lead to large changes in resulting MIND networks (**Supplementary Figure 2**). To empirically select the optimal parameter set for downstream analyses, we anatomically benchmarked networks generated from a range of parameters (**Supplementary Methods**). We chose to vary the percentiles of retained data and number of bins in the histogram algorithm, and the percentiles in the k-NN algorithm as these led to the largest changes in KL estimation (**Supplementary Figure 2**). The optimal parameter set for the histogram algorithm was number of bins = 32 and percentiles = [0.1, 99.9] (**Supplementary Figure 3**). Downstream analyses therefore use this parameter set. Performance of different k-NN parameter networks on anatomical benchmarks was remarkably similar, so we elected to use the default set of percentiles for downstream analyses ([0, 100]).

#### Statistical analysis

*Mean T1w/T2w multiple linear regression covariates*

Some studies undertake additional steps to normalise T1w/T2w scans to improve inter-subject comparability over time. These studies normalise by features such as median white matter intensity (Nerland et al. 2021) and median vitreous humour intensity (Ganzetti, Wenderoth, and Mantini 2014). We conducted sensitivity analyses showing that the inclusion of these features as covariates to our mean T1w/T2w models did not greatly alter the pattern of regional $\beta_{age}$ values (**Supplementary Figure 9B**).

*MIND degree multiple linear regression covariates*

Previous research demonstrated a positive correlation between estimated total intracranial volume (eTIV) and average across all MIND network edges (global MIND; Sebenius et al. 2023), which may reflect allometric scaling relationships between brain size and network connectivity (Ardesch et al. 2022). Though we did not observe a strong positive correlation between eTIV and global MIND (Spearman r = - 0.11), we conducted sensitivity analyses demonstrating that addition of eTIV as a covariate did not greatly influence estimates of regional degree $\beta_{age}$ coefficients (**Supplementary Figure 11C**).

*Spatial autocorrelation-preserving network nulls*

BrainSMASH-generated surrogate T1w/T2w or *MBP* expression maps were used to permute the rows and columns of respective MIND similarity networks, generating null networks with preserved autocorrelation structure. This procedure was applied to the full left hemisphere network then coarse-grained appropriately when axonal connectivity or correlated gene expression matrices were concerned. All statistical tests used 1000 permutations.

#### Age prediction

To assess the extent to which MIND networks capture developmental changes in myelination, we trained machine learning models to predict age from each subject's T1w/T2w scan. The input features (predictors) used were regional mean T1w/T2w, regional weighted degree, and all edge weights. We focused our analyses on animals aged 7 months (pre-pubertal) to 36 months (mature adult). Previous studies have indicated that although the large majority of intracortical myelination occurs before the end of adolescence, increases can continue until mature adulthood (Grydeland et al. 2019). Here, we chose an upper age boundary of 36 months to be reflective of this protracted period of myelination and to ensure that sufficient data were provided for model training, while not extending the upper boundary so far as to increase the chance that models became sensitive to later age-related processes. Furthermore, to ensure our models were not biased by outliers, we removed subjects falling in the top and bottom

1% of mean cortical T1w/T2w or mean weighted degree across all regions. After developmental filtering and quality control, the analysable dataset comprised N=210 subjects (N female = 90).

For each predictor, we trained and tested models on 50 random splits of the data (20% of subjects retained in each test fold), stratified by sex. Within each outer training fold, 5-fold cross validation, again stratified by sex, was used to select between several models. For regional predictors (mean T1w/T2w and weighted degree), we used: a support vector regression with a radial basis function kernel and C regularisation values of [0.1, 1, 10, or 100]; and a gaussian process regression (GPR) with a summed linear and white noise kernel and alpha value of $1 \times 10^{-10}$. We did not use other values of alpha in our final GPR models to reduce computation time and because preliminary tests demonstrated that modulating alpha to [$10^{-5}$, $10^{-3}$, or $10^{-2}$] did not change model predictions. The model with the minimum mean absolute error (MAE) across 5 internal validation folds was trained on the entire training set for that outer fold. At the edge level, due to the much larger feature space (26335 edges) we used only the GPR model (Sebenius et al. 2023).

For each outer test fold, models were evaluated using the mean absolute error (MAE) and the correlation between actual and predicted age values. We used partial correlations to control for the effects of sex and eTIV. Regressing variables out of the feature space can lead to data leakage as estimating effects from the entire dataset allows information from the test set to indirectly influence the training process, inflating model performance (Dinga et al. 2020). We therefore opted to use partial correlations as a post-hoc adjustment (Sebenius et al. 2023).

#### Comparing clustering solutions between modalities

To quantitatively compare T1w/T2w and MBP clustering solutions, we calculated the correlation (Spearman's *rho*) between their cophenetic distance matrices (Sokal and Rohlf 1962). The cophenetic distance between two nodes or cortical areas is defined as the dendrogram height at which they first join the same cluster. The matrix of cophenetic distances between each pair of cortical areas thus summarises the overall structure of the hierarchical clustering solution. Here, high correlations reflect a highly

#### Supplementary Tables

***Supplementary Table 1: Regions used in whole-cortex analysis.*** *Table showing the 115 cortical regions per hemisphere used in our analyses. The amygalo-piriform area of the Paxinos et al. (2012) parcellation was not included in our analyses as it is absent in the BMA 2019 atlas used to parcellate MRI images. Additional anatomical data (cortical zone, cytoarchitectonic class, and position in cortical hierarchy) is listed for all cortical areas for which data is available. Whilst we are using the nomenclature used by Paxinos et al. (2012) for cortical zonal parcellations of marmoset prefrontal cortex, it should be noted that the division of dorsolateral prefrontal cortex extends into regions typically named dorsomedial prefrontal cortex in humans (Carlén 2017).*

| Estimator type | Estimator name | Parameter name | Parameter description | Default parameter value |
|---|---|---|---|---|
| Global | Histogram | Number of bins | Number of bins into which to partition the data | 128 |
| | | Epsilon | Value added to the denominator in Equation (1), when the bin in distribution B is empty, i.e. when $p_b(x) = 0$. This avoids division by 0 errors. | 2.2204e-16 (machine epsilon) |
| | | Percentiles | Percentiles of voxel values to retain. This parameter may be tuned to reduce the influence of outlier values, which increase the range over which bins are stretched and therefore influence the presence of empty bins. | [1, 99] |
| Local | k-NN | *k* | The $k^{th}$ nearest neighbour used to compute local distance in equation (2). | 1 |
| | | Percentiles | Percentiles of voxel values to retain. This parameter may be tuned to reduce the influence of outlier values. | [0, 100] |

***Supplementary Table 2: Tuneable parameters in KL estimators.*** *Tuneable parameters in Histogram and k-NN algorithms, including values used in the default case and a parameter description.*

| Coarse region abbreviation | Coarse region full name | Paxinos subregions (abbreviation) |
|---|---|---|
| DLP | Dorsolateral prefrontal cortex | A10, A46D, A46V, A8aD, A8aV, A8b, A9 |
| VLP | Ventrolateral prefrontal cortex | A45, A47L, A47M, A47O, ProM |
| MPC | Medial prefrontal cortex | A14C, A14R, A32, A32V |
| OFC | Orbitofrontal cortex | A11, A13L, A13M, A13a, A13b, Gu, OPAl, OPro |
| LTC | Lateral temporal cortex | TPO, TPPro |
| LPC | Lateral parietal cortex | OPt, PFG |
| PCA | Posterior cingulate cortex, area 23 | A23V, A23a, A23b, A23c |
| M1 | Primary motor cortex | A4ab, A4c |
| S1 | Primary somatosensory cortex | A1-2, A3a, A3b |
| A1 | Primary auditory cortex | AuA1 |
| V1 | Primary visual cortex | V1 |
| V2 | Secondary visual cortex | V2 |

***Supplementary Table 3: Coarse-grained parcellation used in single cell transcriptomic analyses***. *Coarse-grained cortical parcellation used in single cell transcriptomic analyses and constituent regions. For each of these 12 coarse-grained cortical areas, cells of each subtype were sampled from more than one marmoset, representing both sexes, except for the posterior cingulate cortex, A23, which was sampled in a single male. In total, these cortical data were sampled from 6 marmosets aged from 29 to 34 months, except for one older animal (11 years, 5 months) which contributed one sample from the dorsolateral prefrontal cortex. Given that both sexes were sampled, and predominantly from a constrained age range, transcriptional differences between cortical areas were considered unlikely to be driven mainly by age or sex.*

#### Supplementary Figures

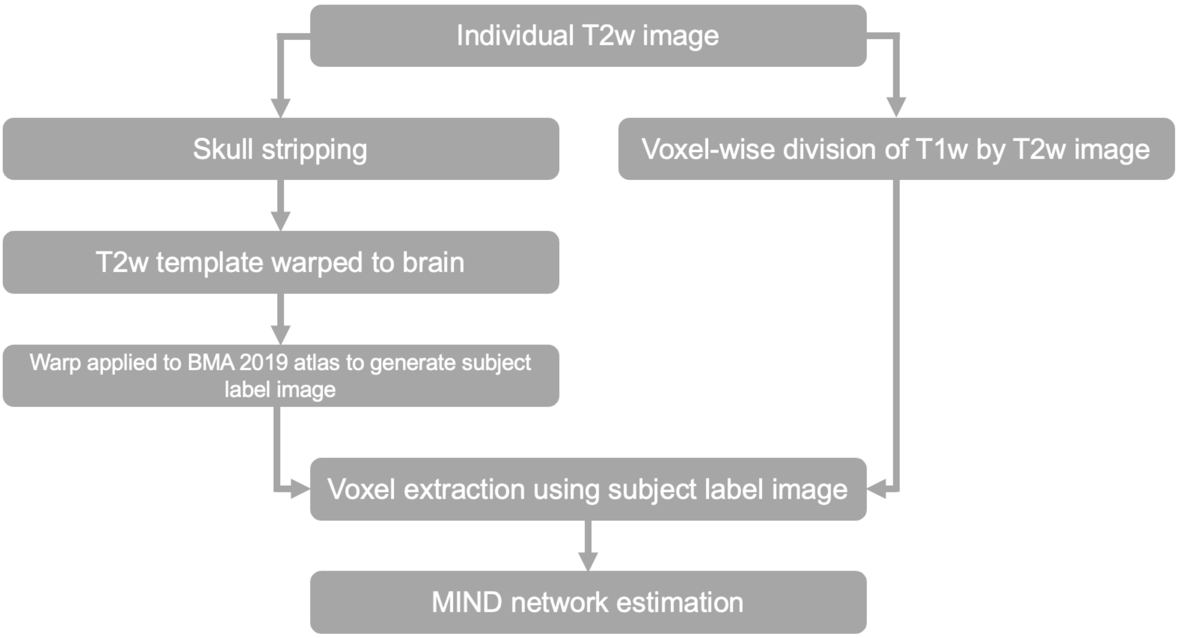


***Supplementary Figure 1: Preprocessing of T1w/T2w ratio images.*** *Schematic showing preprocessing of T1w/T2w ratio MRI images.*

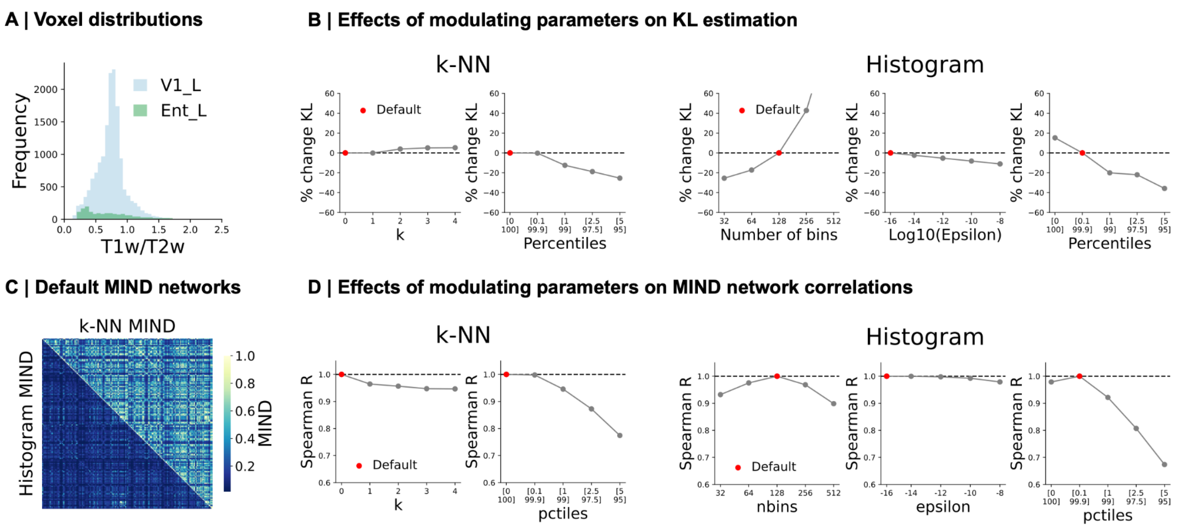


***Supplementary Figure 2: Edge-level effects of tuning parameters in KL divergence estimators. A** Example voxel distributions from an empirical scan (V1_L: left primary visual cortex; Ent_L: left entorhinal cortex). **B** Changing all parameters leads to changes in KL, indicated by the percentage change from default (marked in red). **C** MIND similarity networks estimated with default parameters. **D** Changing all parameters leads to a change in network structure, indicated by a Spearman's rho correlation < 1, from default (marked in red). Changes in **B** and **D** are most notable for k-NN percentiles, Histogram percentiles and number of bins.*

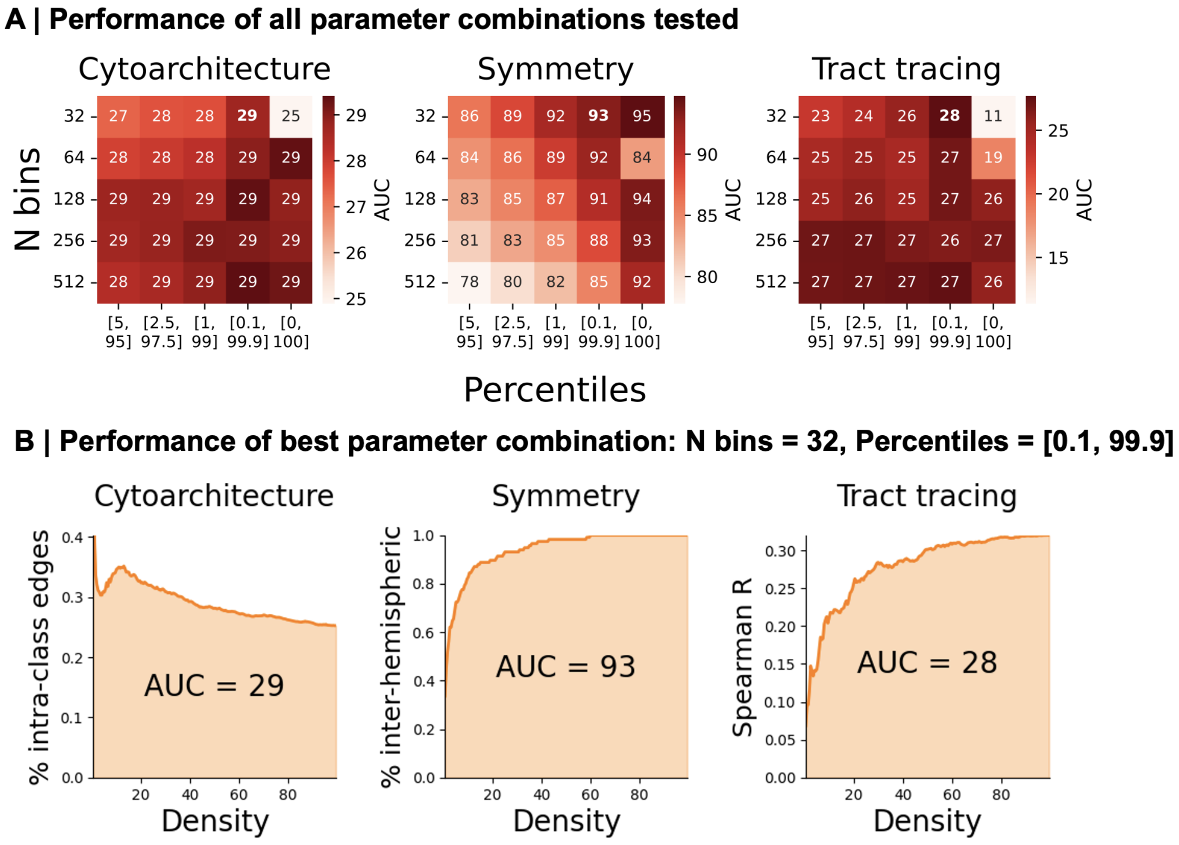


***Supplementary Figure 3: Selecting optimal global algorithm parameters. A*** *Performance of Histogram MIND networks, estimated using all combinations of number of bins and percentiles, on biological benchmarks.* ***B*** *Performance of the optimal Histogram MIND parameter set (Number of bins = 32, percentiles = [0.1, 99.9]), defined as having the highest mean rank performance across all 3 benchmarks in* ***A****.*

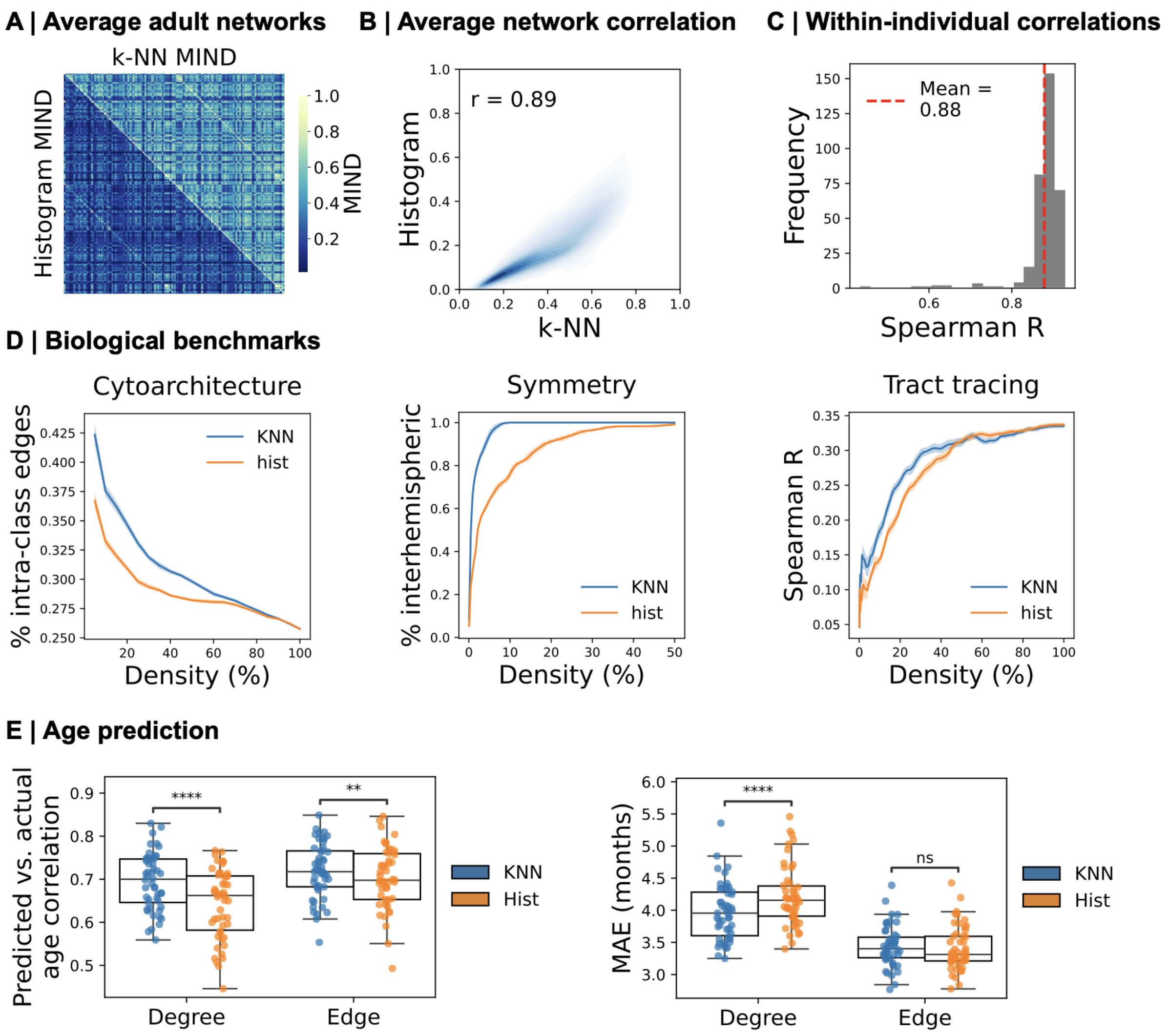


***Supplementary Figure 4 | Local algorithm out-performs global on anatomical benchmarks and age prediction. A** Average adult k-NN MIND and optimised Histogram MIND networks. **B** Average networks in **A** are highly correlated at the edge level (Spearman r = 0.89). **C** Networks of individual subjects are highly correlated at the edge-level (mean Spearman r = 0.88). **D** k-NN MIND out-performs Histogram MIND on cytoarchitectonic, symmetry, and tract tracing benchmarks, indicated by a higher AUC. The shaded areas indicate 95% confidence intervals estimated by subject-level bootstrapping. **E** k-NN MIND out-performs Histogram MIND at age prediction, indicated by a significantly higher median partial correlation between predicted and true age at the degree and edge level, and a significantly lower median MAE at the degree level over 50 splits of the data. Asterisks indicate significance of Wilcoxon test with Benjamini-Hochberg correction for multiple comparisons.*

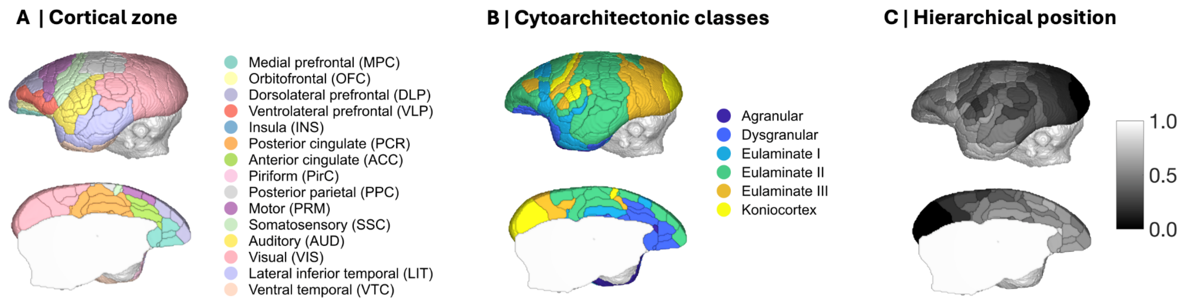


***Supplementary Figure 5: Anatomical maps. A*** *Anatomical and functional cortical zones from Paxinos et al. (2012).* ***B*** *Cytoarchitectonic class assignment map from Atapour et al. (2024).* ***C*** *Hierarchical position map from Theodoni et al. (2021). A full list of cortical areas and anatomical assignments is contained in* ***Supplementary Table 1****.*

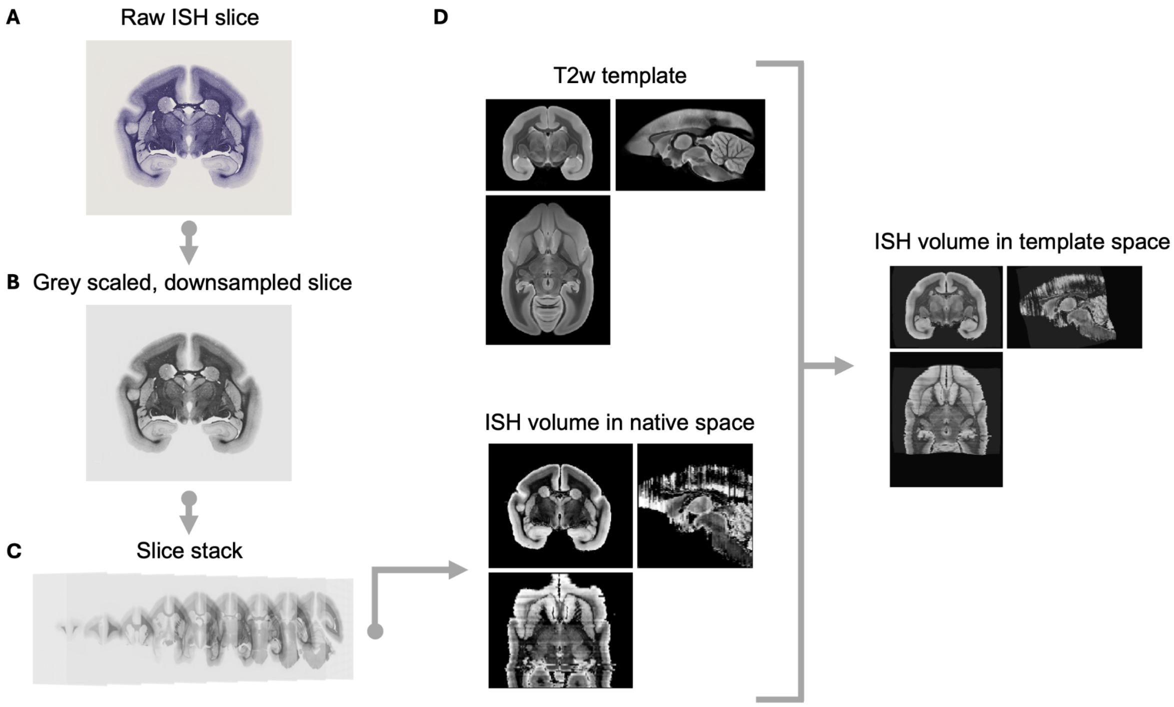


***Supplementary Figure 6: Preprocessing of in-situ hybridisation images.*** *We downloaded in-situ hybridisation images capturing the spatial distribution of myelin basic protein mRNA in a 6 month-old male and 4-year-old male marmoset from the Marmoset Gene Atlas (Shimogori et al. 2018; Kita et al. 2021). In each brain, MBP gene expression was measured in 90 coronal brain slices, which were used to generate an in-situ hybridisation (ISH) volume that was then co-registered to MRI space. Preprocessing of slices used Matlab and ANTs.* ***A*** *Example raw slice.* ***B*** *Full resolution slices were downsampled and gray-scaled to reduce computational load.* ***C*** *Slices were roughly aligned to create a native ISH volume using code from Tong et al. (2022).* ***D*** *The ISH volume was linearly and non-linearly warped to the same ex-vivo T2w template used in the preprocessing of MRI images (Woodward et al. 2018).*

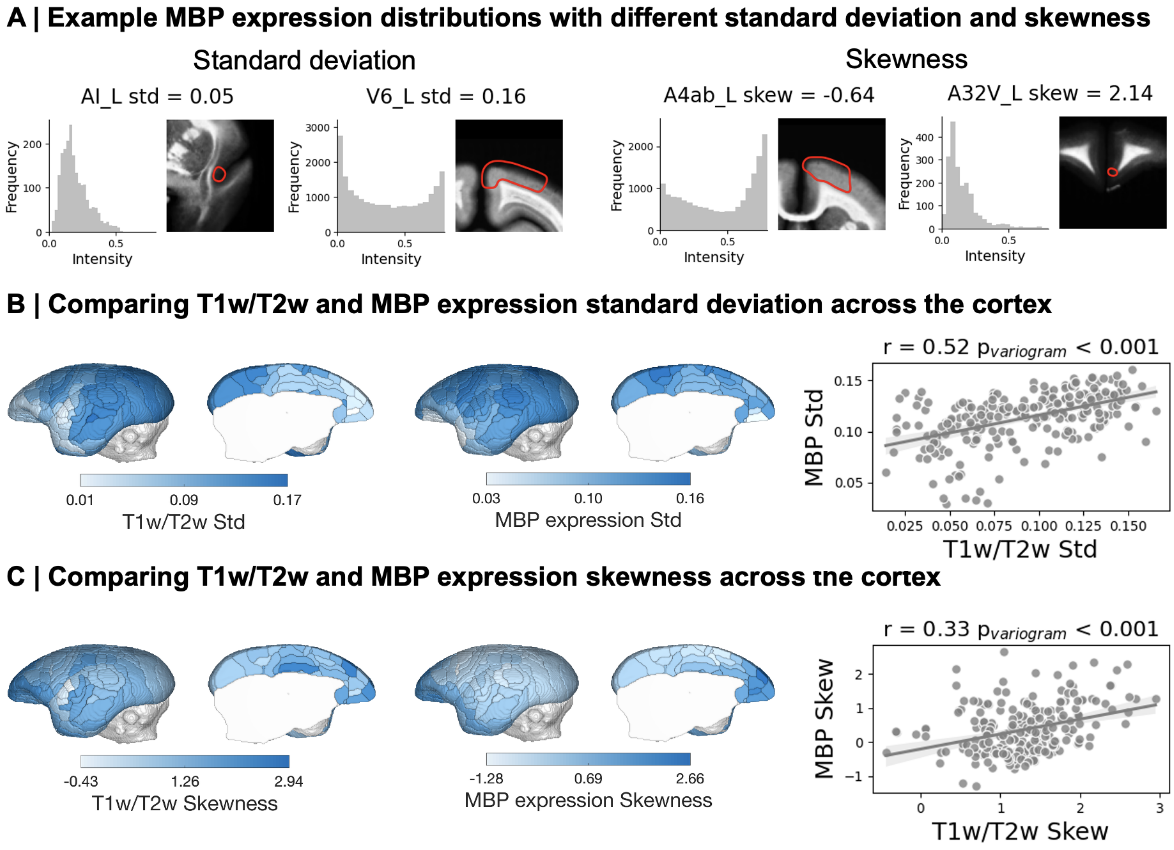


***Supplementary Figure 7: Comparing T1w/T2w and MBP expression distribution shapes. A** Example regions from MBP expression data with low and high standard deviation (left panel) and skewness (right panel). Negative skew indicates that the distribution tail falls to the left, and positive skew to the right. **B** Standard deviations of T1w/T2w and MBP expression distributions are moderately correlated (Spearman r = 0.52, $p_{variogram}$ < 0.001). **C** Skewness of T1w/T2w and MBP expression distributions are weakly correlated (Spearman r = 0.33, $p_{variogram}$ < 0.001).*

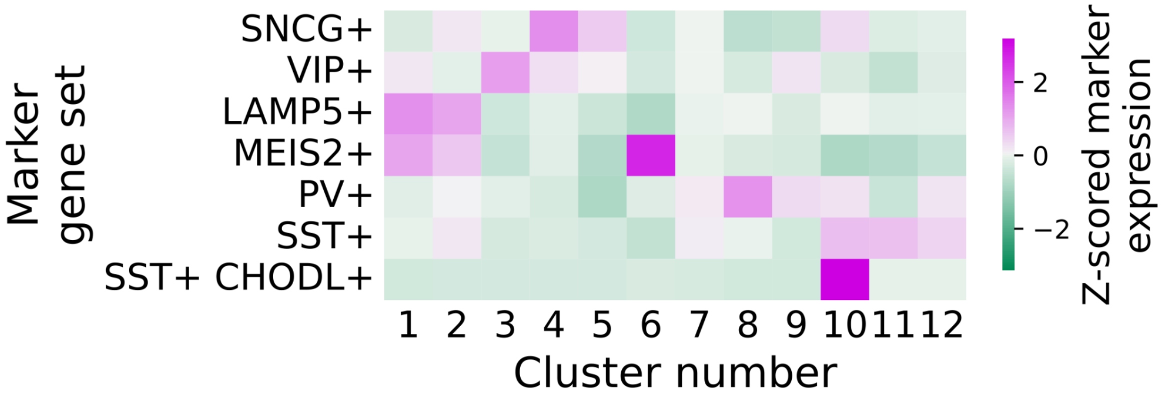


***Supplementary Figure 8: Assignment of interneuron clusters to subtypes**. Marker gene sets generated from marmoset primary motor cortex samples (Bakken et al. 2021) were used to assign interneuron clusters to a biological identity. The mean log expression of each marker gene was computed for each interneuron cluster. Expression of marker genes was standardised by Z-scoring across cell types. The mean Z-scored expression of each marker gene set (shown in the figure panel) was computed for each interneuron cluster, which was used to assign each cluster to a single biological identity. We assigned clusters 1 and 2 to LAMP5+, cluster 3 to VIP+, clusters 4 and 5 to SNCG+, cluster 6 to MEIS2+, clusters 7, 11, and 12 to SST+, clusters 8 and 9 to PV+, and cluster 10 to SST+ CHODL+. Clusters 5 and 6 were excluded from downstream analyses as expression was not present in the 12 cortical areas that formed the focus of our analyses.*

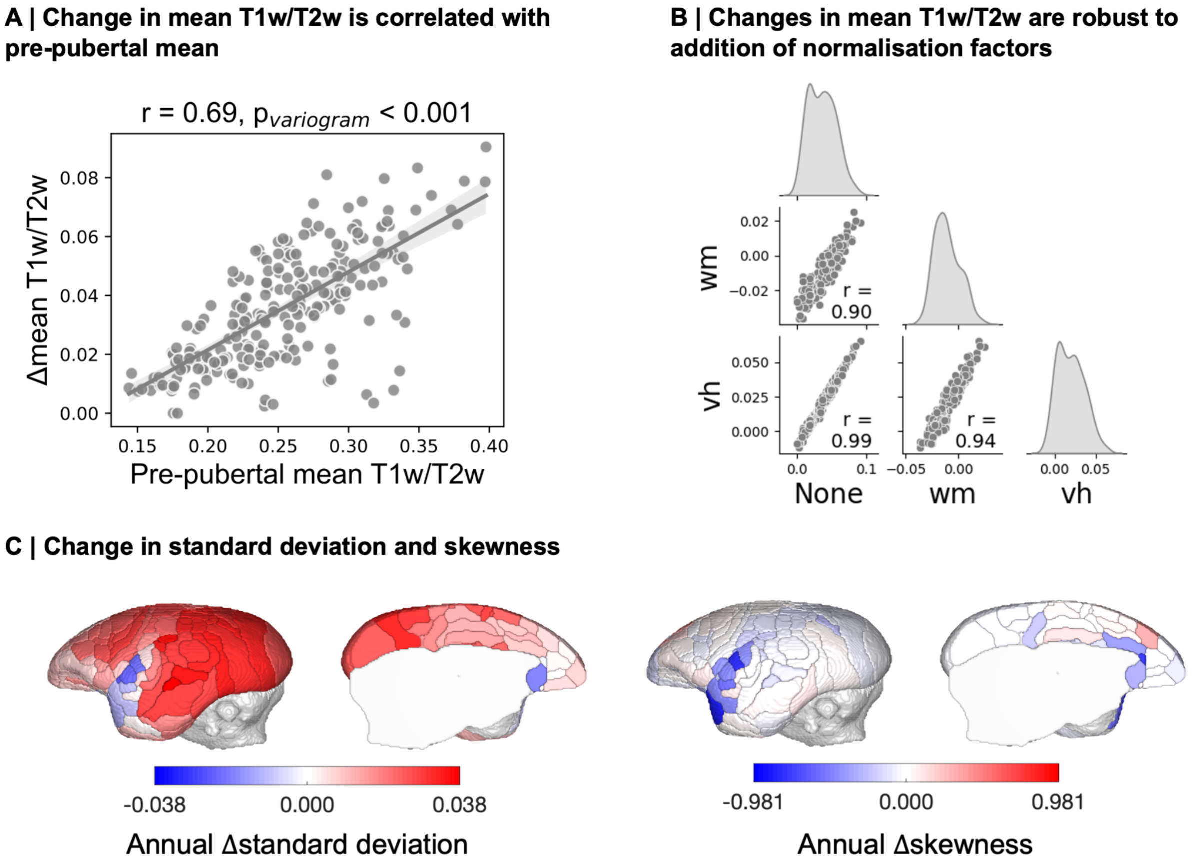


***Supplementary Figure 9: Age-related changes in T1w/T2w. A*** *Pre-pubertal mean T1w/T2w, obtained by averaging across animals <1 year of age, was significantly correlated with a region's adolescent increase in mean T1w/T2w (Spearman r = 0.69,* $p_{variogram}$ *< 0.001).* ***B*** *Sensitivity analysis demonstrating that addition of normalisation covariates to linear models does not greatly alter estimates of age effects on mean T1w/T2w, indicated by high correlations between* $\beta_{age}$ *coefficients for the linear models: Mean T1w/T2w ~ Age + Sex + X, where X is a normalisation covariate: None, median white matter signal (wm) or median vitreous humour signal (vh).* ***C*** *Rate of change in standard deviation (left panel) and skewness (right panel) of T1w/T2w distributions for each cortical area, assessed using multiple linear regressions with age and sex as predictors.*

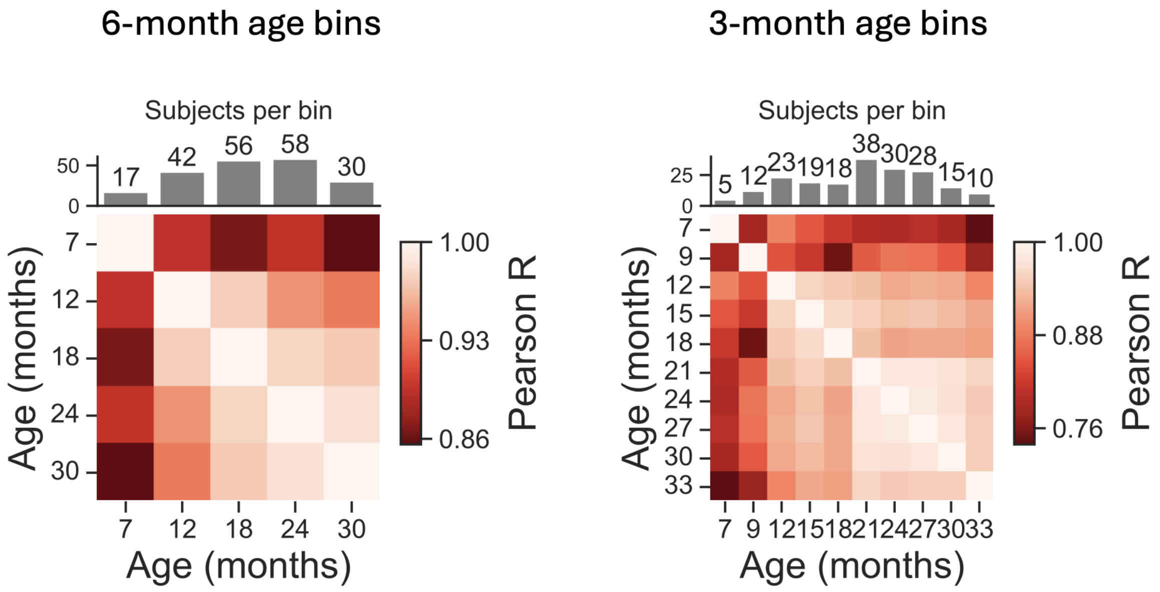


***Supplementary Figure 10: Adult maturational plateau is robust to bin size**. Correlations between age-binned networks remain high after 18 months, when using 6-month age bins, or 21 months, when using 3-month age bins (note different colour bar scales). This indicates that, regardless of bin size, intracortical myelination networks exhibit a slowing of maturational changes close to the onset of adulthood at ~20 months (Sawiak et al. 2018).*

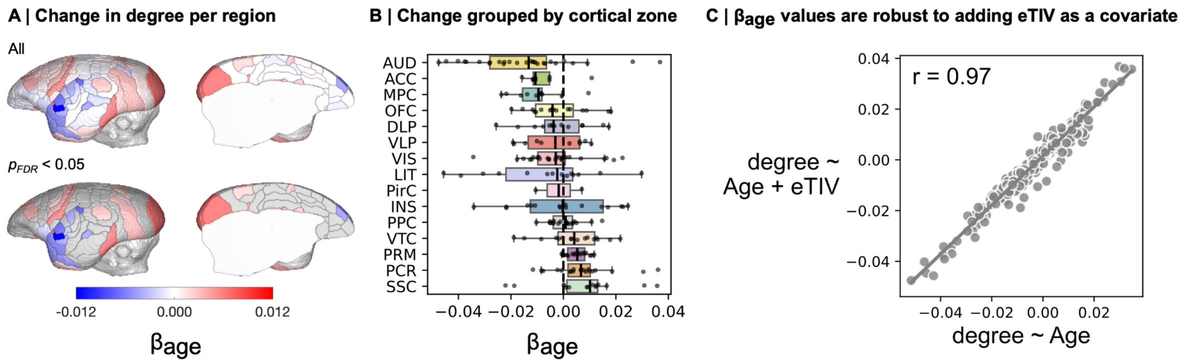


***Supplementary Figure 11: Auditory regions become microstructurally differentiated from the rest of the cortex. A** Multiple linear regressions predicting degree from age and sex. $\beta_{age}$ coefficients for the left hemisphere, uncorrected and corrected for multiple comparisons, are shown. **B** Uncorrected $\beta_{age}$ coefficients grouped by cortical zone. **C** $\beta_{age}$ coefficients are highly correlated when including or excluding estimated total intracranial volume as a covariate.*

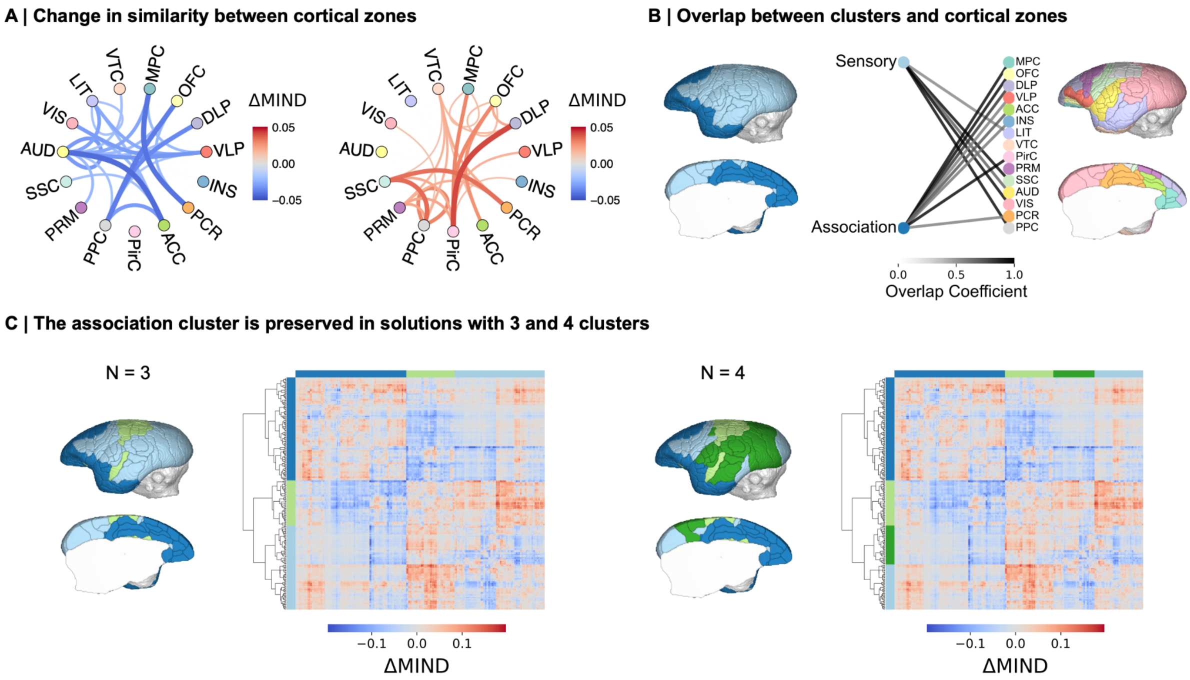


***Supplementary Figure 12: Maturational clustering sensitivity analyses***: ***A*** *Mean rate of change in T1w/T2w MIND between cortical zones of the left hemisphere. The bottom 20% (left) and top 20% (right) of mean changes are shown.* ***B*** *Degree to which cortical zones are contained within each left hemisphere cluster, quantified by the Szymkiewicz–Simpson overlap coefficient. Overlap equals 1 if all regions in a cortical zone are contained entirely within a cluster, and 0 if none of them are. Overlap is shown if greater than 0.5, i.e. if at least half of the areas in a cortical zone are contained within a cluster. The association cluster overlaps principally with frontal and paralimbic association areas, while the sensory cluster overlaps principally with primary sensory and motor areas.* ***C*** *The association maturational cluster is relatively preserved in three and four cluster solutions, suggesting that frontal and paralimbic association areas are particularly strongly maturationally coordinated.*

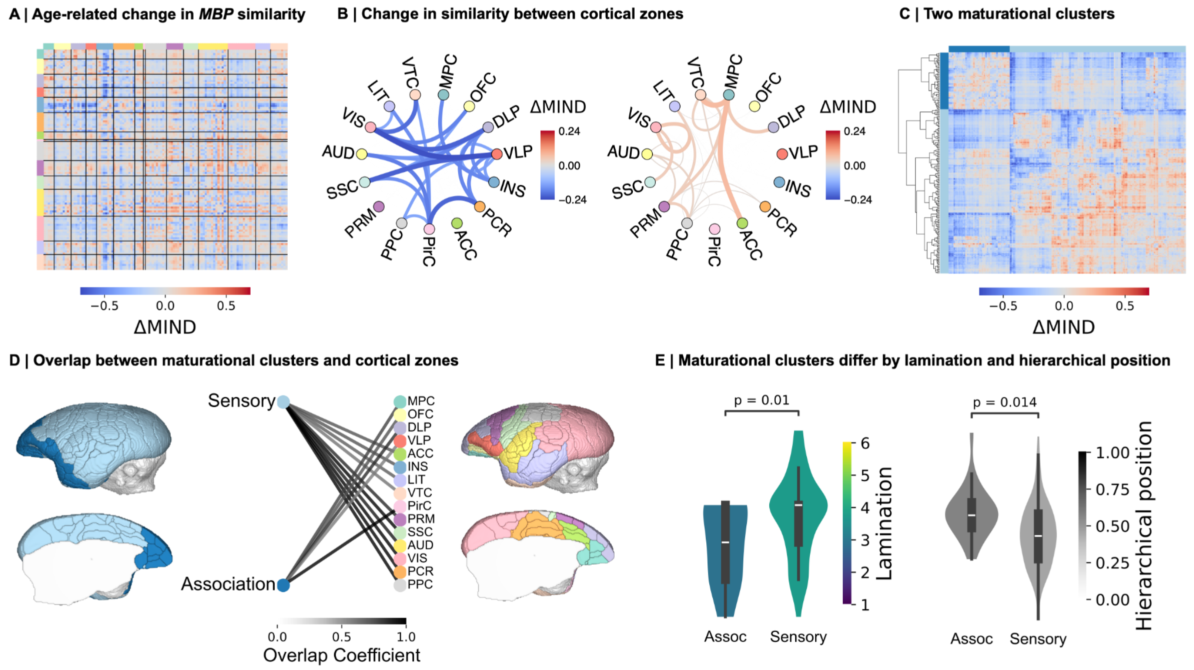


***Supplementary Figure 13: Maturational changes in MBP expression similarity are concordant with changes in T1w/T2w ratio similarity***. ***A*** *The difference between the adult and infant MBP expression MIND network was used to approximate linear change in similarity. Left hemisphere shown, grouped into cortical zones.* ***B*** *Left hemisphere matrix averaged across cortical zones. The bottom 20% (left) and top 20% (right) of mean changes are shown. The divergence between primary sensory (VIS, AUD, SSC) and association (OFC, DLP, VLP, PirC) cortical zones is particularly pronounced.* ***C*** *Hierarchical clustering was performed in an identical manner to the T1w/T2w MIND age-related change matrix.* ***D*** *The degree to which cortical zones were contained within each maturational cluster was quantified using the Szymkiewicz–Simpson overlap coefficient (overlap > 0.5 is shown). MBP MIND development clusters were largely correspondent with T1w/T2w MIND development clusters, though the MBP MIND association cluster contained mainly prefrontal association areas.* ***E*** *Concordant with T1w/T2w MIND development, the sensory cluster had significantly higher mean degree of lamination ($t = 3.72$, $p_{variogram} = 0.010$) and significantly lower mean hierarchical position ($t = -3.69$, $p_{variogram} = 0.014$). Reported p-values were derived from two-tailed, two-sample t-tests evaluated against spatially autocorrelated null models.*
